# Assessing the overlap between fishing activities and chondrichthyans distribution exposes high-risk areas for bycatch of threatened species

**DOI:** 10.1101/2023.10.25.563919

**Authors:** Federico Maioli, Benjamin Weigel, Max Lindmark, Chiara Manfredi, Walter Zupa, Isabella Bitetto, Tommaso Russo, Michele Casini

## Abstract

Sharks, rays, and chimaeras (chondrichthyans) play a crucial role in marine ecosystem functioning but are highly vulnerable to fishing. Hence, understanding the spatial overlap between chondrichthyans and fishing effort is essential for effective conservation and management. Here, we propose an integrated approach that combines Vessel Monitoring System data with geostatistical species distribution models to assess the potential impact of fishing on chondrichthyan populations in the western Adriatic Sea. By mapping the overlap between model-based chondrichthyan distribution, species richness, and the proportion of threathened species with bottom trawl fishing activities, we identify areas at high risk for chondrichthyan bycatch. Our findings show that many of these species are at risk across a large part of their distribution within the study area. Notably, there is a substantial spatial overlap between regions where threatened chondrichthyans are found and species-rich areas with locations of intensive bottom trawl fishing in the northern and central offshore regions of the western Adriatic, emphasizing the vulnerability of these species to fishing pressure. Furthermore, differences in overlap between distinct fishing gears highlight the importance of considering specific fishing practices when formulating management strategies. While our work provides novel insights to potential bycatch hotspots, limitations related to data sources, spatial resolution, and the inability to directly quantify fishing impacts should be considered. Nonetheless, our findings contribute to the development of targeted conservation and spatial management measures, offering a general approach to study model-based spatial hotspots aimed at protecting and sustaining chondrichthyan populations in the heavily exploited Adriatic Sea.

## Introduction

Sharks, rays and chimaeras (chondrichthyans) are crucial for maintaining marine ecosystem functioning (Myers et al., 2007), but at the same time, they are highly vulnerable to fishing due to their slow growth, late maturity and low fecundity (Stevens, 2000). Chondrichthyans are primarily caught as accidental bycatch in fisheries targeting more valuable teleost species, although certain chondrichthyan species are periodically targeted (Bradai et al., 2012; Ferretti & Myers, 2006). The harmful effects of fishing are further exacerbated by environmental threats, including habitat degradation and climate change (Dulvy et al., 2014, 2021). These combined factors have led to a widespread decline in chondrichthyan populations over the past few decades (Pacoureau et al., 2021; Simpfendorfer et al., 2023) and have placed many species at risk of extinction (Dulvy et al., 2021). Such declines could, in turn, have adverse impacts on marine ecosystems (Ferretti et al., 2010).

Spatial management of fisheries is nowadays a largely-applied approach to mitigate the adverse impacts of fisheries (FAO, 2023; McConnaughey et al., 2020). In this context, assessing the spatial overlap between chondrichthyans and fishing effort is crucial for understanding the potential impacts of fishing on these vulnerable species, and thus for designing effective conservation and management strategies (Jorgensen, 2022; Queiroz et al., 2016, 2019). However, limited data on incidental catches, landings, and species identification in many regions hinder the understanding of where, when, and which chondrichthyan species and fishing vessels overlap across their ranges, thereby limiting the implementation of targeted management measures (Cashion et al., 2019; FAO, 2022).

In cases where direct data, such as logbooks, offering accurate and reliable information about catches of these species are unavailable, a widely adopted approach involves the use of tracking devices to monitor the movements of fishing fleets in both space and time (Bastardie et al., 2010; Russo et al., 2014). Specifically, Vessel Monitoring Systems (VMS) have emerged as a valuable tool for tracking and monitoring fishing effort (Amoroso et al., 2018; Eigaard et al., 2017). Through satellite-based communication systems, VMS allows for real-time tracking of fishing vessels, thus facilitating the identification of areas with high fishing activity (Witt & Godley, 2007). By integrating VMS data with species distribution information, a more comprehensive understanding of the spatial overlap between fish and trawling activities can be achieved. This integrated approach may help identify areas of high risk for chondrichthyans bycatch and therefore guides targeted conservation efforts.

The Adriatic Sea, situated in the central Mediterranean, has a long history of chondrichthyans fisheries and has experienced depletion of several species in the last few decades (Dulvy et al., 2003; Ferretti et al., 2013; Jukic-Peladic et al., 2001; Lotze et al., 2011). Among the chondrichthyan species commonly caught or observed in this region are the dogfish (*Scyliorhinus* spp.), brown skate (*Raja miraletus*), spurdog (*Squalus acanthias*), thornback skate (*Raja clavata*), and smooth-hound (*Mustelus* spp.) (Barausse et al., 2014; Clodia database, 2020; Ferretti et al., 2013; Maioli et al., 2023). Presently, approximately 70% of the chondrichthyan species in the Adriatic Sea are regionally threatened according to International Union for Conservation of Nature (IUCN) Red List Criteria (www.redlist.org) (Soldo & Lipej, 2022). This basin is recognized as one of the most trawled regions globally (Amoroso et al., 2018; Pitcher et al., 2022), with bottom trawling being a significant and widely practiced fishing activity targeting various demersal fish species (FAO, 2022). The bottom trawling fleet in the Adriatic Sea consists of 1946 vessels, equivalent to approximately 64,900 gross tonnes, with Italy owning around 70% of the total fleet (FAO, 2022). This fishing activity significantly contributes to the incidental capture of chondrichthyans, making it a major concern for the conservation of these species (FAO, 2022). Recognizing the need for action, the General Fisheries Commission for the Mediterranean (GFCM) initiated the MedBycatch project. Its goal is to develop a collaborative approach to improve the understanding of the Mediterranean multi-taxa bycatch of vulnerable species and test mitigation actions (Carpentieri et al., 2021). However, despite these efforts, there is limited knowledge regarding the bycatch of chondrichthyan species, which hampers effective management and conservation efforts in the region (FAO, 2022).

In this study, we adopt a novel approach to examine the spatial co-occurrence between chondrichthyan species and commercial fishing practices using the Adriatic Sea as a case study. We integrate data on fishing effort from the Italian fleets obtained through VMS with predicted species distributions derived from geostatistical Species Distribution Models (SDMs) fitted to data from a fishery-independent bottom trawl survey. Additionally, we analyze and map species richness and the proportion of threatened species. This enables us to assess the overlap between this community and bottom trawl fishing activities, providing insights into the potential impact of fishing. Our goal is to identify high risk areas for bycatch, where the potential impact of fishing on chondrichthyan populations and communities is the highest, thus guiding the development of targeted conservation efforts and management strategies to protect chondrichthyan populations in this heavily exploited ecosystem.

## Methods

### Study area

The Adriatic Sea, located in the Central Mediterranean between the Italian peninsula and the Balkans, is primarily a shallow and eutrophic basin (Appendix S1). It displays distinct morphological variations along both its longitudinal and latitudinal axes.

In the northern part of the Adriatic Sea, there is an extremely shallow mean depth of approximately 30 m up to a maximum of 70 m, and the bathymetric gradient along the major axis of the basin is quite weak (Appendix S1). This region experiences a strong river runoff, with the Po River and other northern Italian rivers contributing around 20% of the entire Mediterranean river outflow (Hopkins, 1992). These rivers introduce significant amounts of nutrients, making the Northern Adriatic Sea the most productive area in the Mediterranean (Campanelli et al., 2011) and one of the most heavily fished regions in Europe (Eigaard et al., 2017).

The central zone of the Adriatic Sea reaches depths of about 200m. However, it is also characterized by two depressions known as the Jabuka/Pomo Pits, which have a maximum depth of approximately 270 m.

The Southern Adriatic Sea exhibits significant differences between its northern and southern areas. In the northern region, specifically around the Gulf of Manfredonia, a wide continental shelf can be found, with a distance of about 45 nautical miles between the coast and the 200-meter isobath. This area is characterized by a smooth slope. On the other hand, in the southern region, the isobaths are much closer together, with a distance of only 8 nautical miles. These distinct morphological variations between the two areas have direct implications for the biocenoses, fisheries resources, and prevalent fishing techniques employed in the region (Anonymous, 2011).

The western coast of the Adriatic Sea is generally regular, sandy, and gently sloping, while the eastern coast is characterized by irregularities, numerous islands, and steeply sloping rocky bathymetry (Russo & Artegiani, 1996).

The water circulation pattern in the Adriatic Sea is cyclonic, with water masses flowing into the basin from the Eastern Mediterranean following a north-westward flow on the east oat and exiting trough a south-eastward flow on the west coast. The circulation and water masses in the Adriatic Sea are strongly influenced by river runoff and atmospheric conditions, which, in turn, impact the salinity and temperature of the water (Artegiani et al., 1997).

### Fishing effort data

Vessel Monitoring Systems (VMS) data consists of a series of consecutive pings (signals) sent by each vessel at regular time intervals. Compared with other tracking devices like the Automatic Identification System (AIS), the temporal frequency of these pings is relatively low, usually with 1-2 hours between successive pings. However, the VMS data offers unparalleled spatial coverage as it relies on the INMARSAT satellite network (Russo et al., 2016; Shepperson et al., 2018). The main limitation of VMS is related to its limited coverage of the fleets since VMS is mandatory only for fishing vessel with a length-over-all larger than 12 m, which means that smaller vessels may not be included in the data.

For this study, VMS and logbook data for the trawling fleet operating in the Adriatic Sea were provided by the Italian Ministry of Agriculture, Food Sovereignty and Forests (MASAF) within the scientific activities related to the Italian National Program for the Data Collection in the Fisheries Sector (INPDCF). We extracted a subset of VMS data from 538 vessels utilizing bottom otter trawl (OTB) and beam trawl (TBB) gear types, which are two common fishing methods used along the Italian Adriatic coast. These bottom trawlers were identified through cross-analysis involving logbook data and the EU Common Fleet Register (https://webgate.ec.europa.eu/fleet-europa/index_en).

This subset of data was then used to assess the fishing activity of these vessels from 2009 to 2021 using the R package VMSbase (version 2.2.1; Russo et al., 2014; Russo et al., 2011a; Russo et al., 2011b). VMS pings were partitioned into fishing trips by vessel and interpolated to increase their temporal frequency to 10 minutes (Russo et al., 2011a). The high-frequency interpolated VMS pings were then joined with the NOAA-Etopo1 database (Amante, Christopher & Eakins, 2009) through the R package marmap (Pante & Simon-Bouhet, 2013) to estimate the bottom sea depth for each ping. Finally, to identify fishing activities accurately, fishing set positions were isolated from other vessel states, such as steaming and resting, by applying a combination of speed and depth filters (Russo et al., 2014).

Thereafter, we filtered the data to include only the period covered by the MEDITS survey (May to September, see below). Subsequently, we computed the yearly aggregated value of trawl fishing effort (i.e. the sum of the fishing hours) for each cell in a 2 × 2 km square grid multiplying the number of fishing set positions by the interpolation frequency (10 minutes). We then filtered this grid by excluding depths not covered by the MEDITS survey, specifically excluding depths < 10 m and > 800 m. Additionaly we excluded the national territorial seas of Slovenia, Croatia, Montenegro and Albania. The shapefiles for the exclusion of the territorial seas was obtained from www.marineregions.org (2023).

### Survey data

The complete biological dataset considered for this study consisted of the catches of 5,122 bottom trawl hauls that were performed in the Italian and international waters of the Adriatic Sea during the Mediterranean International Trawl Survey (MEDITS). This survey has been conducted annually since 1994 by the Laboratory of Marine Biology and Fisheries of Fano (Italy) in the Northern and Central Adriatic Sea and by Laboratorio Provinciale di Biologia Marina of Bari (Italy) (1994-2008) and COISPA Tecnologia & Ricerca (since 2009) in the Southern Adriatic Sea. The survey is generally carried out in the late spring-summer period (May to September, although in some years it has been performed in October-December), and the hauls are located at depths ranging from 10 to 800 m, following a random-stratified sampling scheme based on 5 different depth strata (Anonymous, 2017; Spedicato et al., 2020). The GOC-73 experimental bottom trawl is used as the sampling gear, which has a horizontal opening of 16-22 m and a vertical opening of approximately 2.4 m. The codend of the trawl is equipped with a 20 mm side diamond stretched mesh. Further information on the sampling procedures, data collection, and analysis can be found in the MEDITS handbook (Anonymous, 2017).

We selected and analyzed a total of 4,216 hauls in this study (Appendix S1), after excluding hauls conducted prior to 1999 and hauls conducted during the autumn period (October to December). The decision to exclude these hauls was driven by two factors. Firstly, the availability of the biogeochemicals variables (see section *Environmental covariates and model selection*) only reaches back to 1999. Secondly, the coverage of the autumn season has been inconsistent over the years. Ensuring a seasonally homogenous dataset was essential, considering the potential redistribution of chondrichthyan species across seasons (Manfredi et al., 2010). Moreover, we excluded species scarcely represented in the data (i.e., those that occurred in less than 3% of all the trawl hauls considered). Further, we grouped the common smooth-hound (*Mustelus mustelus*) and the blackspotted smooth-hound (*Mustelus punctulatus*) together under the category smooth-hound (*Mustelus* spp.) because the morphological identification for the *Mustelus* genus is not straightforward (Marino et al., 2018), and the two species can therefore be misclassified. Additionally, the two species occur together in the Northern Adriatic Sea, display a similar diet (Di Lorenzo et al., 2020), inhabit similar habitats, and the chance of hybridization is not excluded (Marino et al., 2015, 2018). In total, 10 species/genera were included in further analyses (Table 1).

**Table 1.**
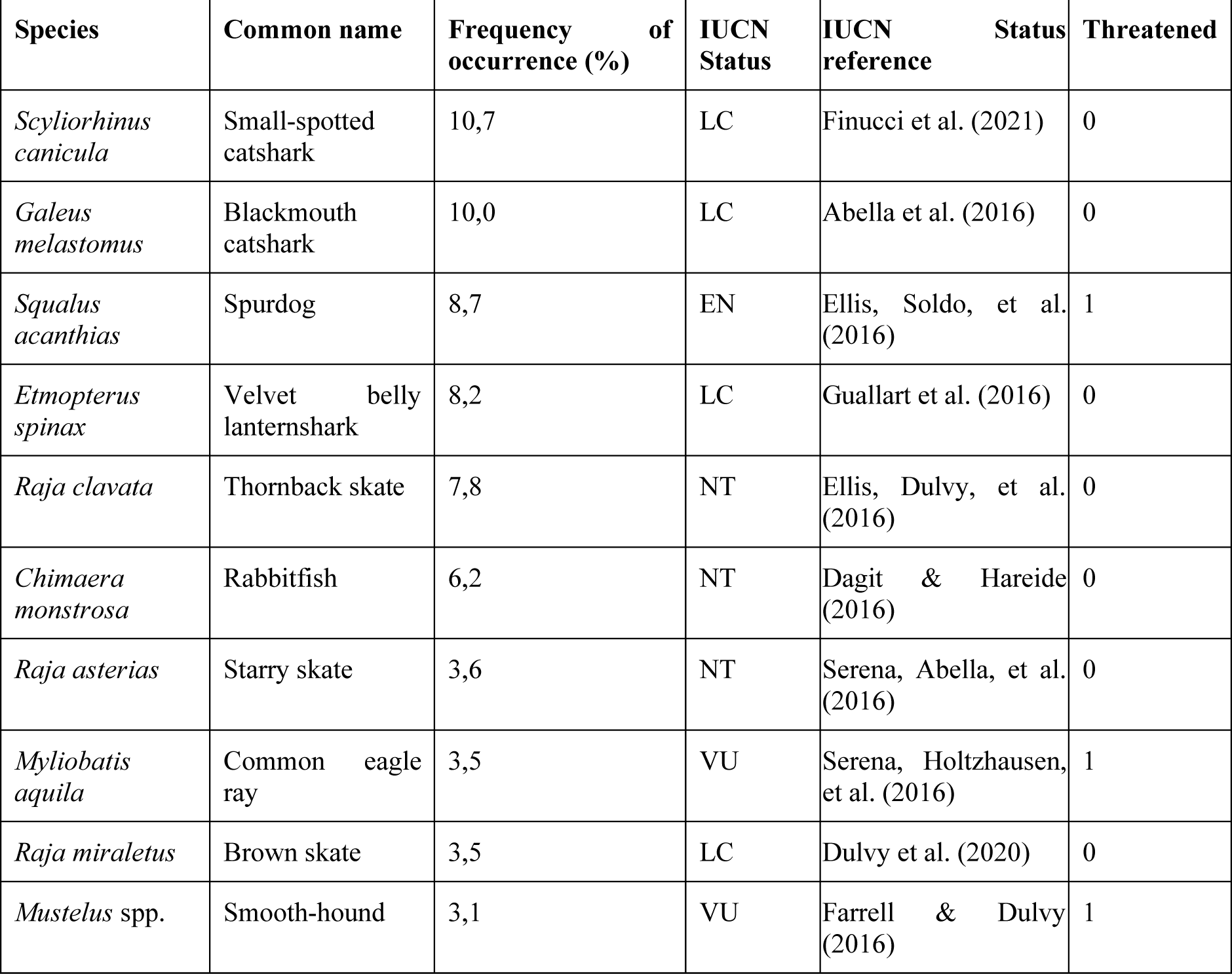
Species/genera selected for this study and the corresponding frequency of occurrence (percentage of trawl hauls where the taxon was caught) for the period 1994-2021. Species are sorted in descending order of occurrence. The IUCN Status column indicates the conservation status of each species: EN (Endangered), VU (Vulnerable), NT (Near Threatened), and LC (Least Concern). In this study, species are categorized as nonthreatened (0) if classified as LC or NT and threatened (1) if classified as EN or VU.

### Species distribution models

We utilized geostatistical generalized linear mixed-effect models (GLMMs) as SDMs to capture the distribution patterns of chondrichthyan species (Table 1). These models combined survey data and environmental covariates, while also considering the influence of spatial and spatiotemporal correlations using Gaussian random fields.

Specifically, we modelled the presence/absence of each species using a Bernoulli distribution:

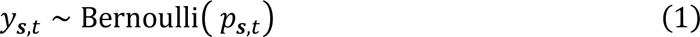

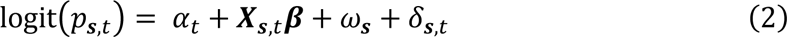

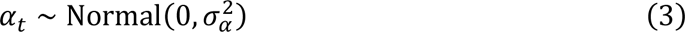

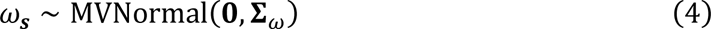

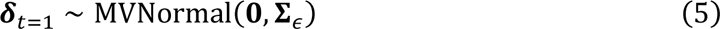

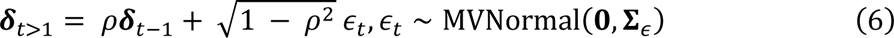

where *y*_***s**,t*_ is the binary response variable (presence/absence) for the observation at space ***s*** and year *t*, *p*_***s**,t*_ denotes the probability of observing the species at space ***s*** and year *t*, *α*_*t*_ represents random intercepts by year *t*. ***X******_s_****_,t_* represents the vector of the environmental covariates (see the section *Environmental covariates and model selection* and Appendix S2) at space ***s*** and year *t* and *β* represents a vector of corresponding coefficients. ω_***s***_ and *ϵ*_***s**,t*_ are the spatial and spatiotemporal random fields with estimated covariance matrices Σ_ω_ and Σ_ɛ_, respectively. Spatiotemporal random effects are here assumed to follow a stationary AR1-process where *ρ* represents the correlation between subsequent spatiotemporal random fields. To overcome convergence issues encountered during the initial analysis, for the deep-sea species (blackmouth catshark, rabbitfish, and velvet belly lanternshark) and the rare species (with a frequency of occurrence ≤ 5%: starry skate, common eagle ray, brown skate, smooth-hound) we did not include spatiotemporal random fields (*ϵ*_***s**,t*_) in the models and relied exclusively on the spatial random fields (*ω*_***s***_).

To optimize computational efficiency, we utilized a predictive process modeling framework (Anderson & Ward, 2019; Latimer et al., 2009) for fitting all models. In this framework, we employed the Stochastic Partial Differential Equation (SPDE) approximation (Lindgren et al., 2011) and a triangulated mesh (Appendix S3), generated using the R-package R-INLA (Rue et al., 2009), to represent spatial random fields. Random effects are estimated at the knots (the vertices of the mesh) and are then bilinearly interpolated to data locations. The optimal knot locations were determined using a *k*-means clustering algorithm, which minimized the total distance between knots and data points. We selected 200 knots for our models after an initial exploration, as this balance between accuracy and computational time was deemed appropriate (Appendix S3). The models were fitted using TMB (Kristensen et al., 2016) through the R-package sdmTMB (version 0.2.1) (Anderson et al., 2022 [preprint]) with maximum marginal likelihood and the Laplace approximation to integrate over random effects. We assessed convergence of the models and ensured that the maximum absolute gradient with respect to all fixed effects was < 0.001, and that the Hessian matrix was positive-definite (Anderson et al., 2022 [preprint]). The model residuals are presented in the Appendix S4-S7.

### Environmental covariates and model selection

For the SDMs, we selected environmental covariates based on data availability and their assumed relevance to the distribution of the studied species (Maioli et al., 2023). We included four covariates: depth (m), seafloor temperature (°C; hereafter temperature), seafloor dissolved oxygen (ml/L; hereafter oxygen), and seabed substrate (categorized into “Sandy mud”, “Fine mud”, “Sand” and “Muddy sand”). We extracted monthly mean temperature and oxygen values from the Copernicus Marine Service (Escudier et al., 2020; Teruzzi et al., 2021), wich have a spatial resolution of approximately 4 km. We obtained seabed substrate and depth data from the EMODnet Seabed Habitats project (https://www.emodnet-seabedhabitats.eu) and the EMODnet Bathymetry project (https://www.emodnet.eu/en/bathymetry), respectively. To integrate these environmental covariates with the trawl hauls data, we associated each trawl haul with the nearest spatial point of the variables. In the case of temperature and oxygen variables, we also matched the trawl hauls data with the nearest available measurements in time. Inter-annual variations were accounted for by including year as random intercept.

We then asked which combination of covariates resulted in the most parsimonious model, as estimated by the Akaike Information Criterion (AIC). To investigate the relationship with depth, for the most common species (spurdog, small-spotted catshark, and thornback skate), we explored two representations to account for bell-shaped relationships: log(depth) + log(depth^2^), and depth + depth^2^. Additionally, we evaluated whether a linear term alone or a combination of linear and quadratic terms for temperature provided a better fit to the data. Finally, we tested the inclusion of substrate type and of the interaction term between oxygen and temperature.

To address convergence issues encountered during the initial analysis, we narrowed down our model selection process for the deep sea species and for the rare species. For the deep sea species we focused on the functional form of the depth covariate. For the rare species, we focused on the functional forms of both depth and temperature covariates. Specifically, for depth, we considered the following options for both deep sea species and rare species: depth, log(depth), depth + depth^2^, and log(depth) + log(depth^2^) (Appendix S2).

Moreover, for the small-spotted catshark, the velvet belly lanternshark and the thornback skate, we refitted the best candidate model excluding the random year intercepts, as their effect sizes were small and yielded convergence issues. For the common eagle ray we set Normal(0,10) weakly informative priors for the linear and quadratic terms of the depth covariate, as the initial estimates were unreasonably high, and consequently, we refitted the best candidate model using a penalized likelihood framework.

In total, we compared 16 different models for each common species, 4 for each deep sea species, and 8 for each rare species. To facilitate comparison of effect sizes between the included covariates and to compare them to the marginal standard deviation of spatial variation, we rescaled all covariates to have a mean of 0 and a standard deviation of 1 (Schielzeth, 2010). Finally, the explanatory power of the models was evaluated by computing the R^2^ values (Nakagawa et al., 2017; Nakagawa & Schielzeth, 2013) and the Area Under the Curve (AUC; Pearce & Ferrier, 2000).

### Prediction grid

We predicted the probability (*p*) of chondrichthyan species occurrence onto the same 2 × 2 km grid used for the assessment of the trawl fishing effort, using the best-fitting models. We followed the previously mentioned methodology by matching environmental covariates to the midpoint of each cell. For this particular step, we utilized the average values of temperature and oxygen spanning the core survey period, which extends from May to September. By considering the mean values over this period for each cell, we standardized the effects of temperature and oxygen, making our predictions comparable across different years. We restricted the predictions to the period between 2009 and 2021, as VMS data were unavailable for earlier years.

### Species richness

To estimate the species richness, we predicted onto the grid by drawing 1000 simulations from the joint precision matrix of the best-fitting model for each species. This allowed us to obtain an estimate of species richness by summing the number of species in each grid cell (Ovaskainen & Abrego, 2020).

### Proportion of threatened species

To characterize the chondrichthyan community as a whole, we calculated the proportion of threatened species. This was done for each cell of the prediction grid, weighing the species-specific conservation status by the predicted proportion of each species in each cell. To determine the conservation status of each species, we used the criteria set forth by the IUCN (www.redlist.org), categorizing species as threatened (1) if they were classified as Critically Endangered (CR), Endangered (EN), or Vulnerable (VU), and as nonthreatened (0) if they were categorized as Near Threatened (NT) or Least Concern (LC) (Table 1). Spurdog, common eagle ray and smooth-hound are categorized as threatened species, whereas the remaining species are classified as nonthreatened (Table 1).

### Spatial Overlap Index

To evaluate the relationship between the distribution of chondrichthyans and fishing activities, we developed a species-specific Spatial Overlap Index (SOI).

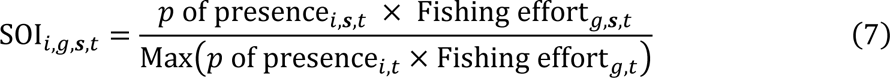

The calculation for the SOI involves multiplying the predicted probability (*p*) of species *i* being present in a specific grid cell ***s*** for a given year *t* by the corresponding fishing effort (measured in fishing hours) for gear type *g* during the same year *t* and grid cell ***s***. The resulting values are subsequently divided by the maximum value obtained from the product of predicted probabilities with fishing effort for gear type *g* and year *t*. This rescaling ensures interpretability and constrains the index within a range of [0,1], and puts a stronger emphasis on the spatial overlap patterns. Consequently, we presented the SOI as an average across the most recent years (2018-2021) since we consider this timeframe more relevant from a conservation point of view. Also, since 2018 a Fisheries Restricted Area was established in the Jabuka/Pomo Pits banning demersal fisheries (GFCM, 2021).

### Range overlap

We employed a range overlap metric (RO) as defined in Carroll et al. (2019) to quantify the overlap between chondrichthyan species and trawling activities. The RO metric estimates the proportion of a species’ area of occupancy where also trawling activities occur:

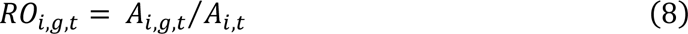

Where *A*_*i,g,t*_ is the area occupied by the species *i* and trawled by gear *g* in year *t*, while *A*_*i,t*_ is the total area occupied by species *i* in year *t* (Saraux et al., 2014). Here, occupied cells were defined as cells with ≥ 75th percentile of the species maximum probability of occurrence. Similarly, to identify cells with high trawling activities, we used a threshold where the fishing effort corresponds to ≥ 75th percentile of the maximum fishing effort for a gear type. Here, we normalized yearly fishing effort within the range [0,1]. This normalization helps address the potential limitation of VMS data as an accurate proxy for overall fishing effort. Due to the pronounced zero-inflation in grid-cell level VMS data, caused by the spatially heterogeneous fishing effort, we calculate percentiles based only on cells with fishing effort > 0. Cells with a fishing effort of 0 are therefore assigned a value of 0. While an alternative approach involves computing percentiles across the entire domain and selecting a higher threshold, we find it more intuitive to interpret the percentiles of presences in this zero-inflated scenario. To ensure the robustness of the chosen threshold, we also explored the sensitivity to different set thresholds.

### Species richness and proportion of threatened species overlap with fishing effort

Finally, we explored the chondrichthyans community overlap with fishing effort, using the estimated species richness and the proportion of threatened species. To visualize these relationships, we employed the R-package biscale (Prener et al., 2022) to generate bivariate choropleth maps. To this end, we used fishing effort, species richness, and the the proportion of threatened species as averages across the most recent years (2018-2021).

## Results

### Fishing effort

Overall, we processed an average of 445 unique VMS identifiers for otter bottom trawlers (OTB) and 93 for beam trawlers (TBB) per year (Appendix S8). Otter trawling was extensively practiced throughout the entire region (Appendix S9). On the other hand, beam trawling was primarily concentrated in the northern regions, extending along the coastline towards the central part of Italy (Appendix S9). Fishing effort (fishing hours in our study) was substantially higher for OTB compared with TBB. The temporal variation over space of OTB was relatively lower (lower coefficient of variation, CV) compared with TBB (Appendix S9), which can be attributed to the lower fleet size of the latter (Appendix S8).

### Species distribution models

Considering the entire survey period (1999-2021), the most dominant species were small-spotted catshark and blackmouth catshark being observed in ∼ 10% of the hauls, followed by spurdog, velvet belly lanternshark, thornback skate, and rabbitfish being recorded in ∼ 8% to 6% of the hauls, while starry skate, common eagle ray, brown skate, and smooth-hound occurred more sporadically (Table 1).

All species models demonstrated a good fit to the data, with conditional R^2^ values ranging from 0.54 to 0.93, and AUC values ranging from 0.91 to 0.99 (Appendix S2). Overall, the inclusion of the linear and quadratic terms for depth provided better support to the models (i.e. lower AIC) across species. Specifically, half of the species models showed a better fit using log-transformed depth while the other half performed better without the transformation (Appendix S2). 3 species models exhibited better fitting when using the linear temperature term, another 3 benefited from employing both the linear and quadratic termperature terms, and for 1 species model, the inclusion of temperature was not supported. Among the 3 species for which the interaction term between temperature and oxygen was considered, the inclusion of this term led to better model fitting in 2 cases (Appendix S2).

The models’ predictions revealed a distinct spatial pattern in the chondrichthyan community (Appendix S10, S11). Specifically, among the threatened species, spurdog exhibited an elevated average probability of occurrence (*p*) across the northern and central offshore zones. Similarly, common eagle ray and smooth-hound displayed higher average *p* values in the northern coastal regions, extending into the offshore area below (Appendix S10).

Conversely, for nonthreatened species, the small spotted catshark, thornback skate, and brown skate predominantly occupied the central offshore areas and the southeastern part of the study area (Appendix S11). Deep-sea species such as the blackmouth catshark, velvet belly lantern shark, and rabbitfish were prevalent in the deeper southern central regions. The starry skate, on the other hand, showed a more prevalent distribution along the central and southern coastal zones (Appendix S11).

Species richness exhibited a distinct contrast between the coastal central and southern regions, which were relatively poor in species, and the northern and deeper southern regions, which were rich in species (Fig. 1a). In the northern areas, the relatively high probability of occurrence of spurdog, smooth-hound, and common eagle ray (Appendix S10) contributed to the observed higher species richness. On the other hand, the higher species richness observed in the deeper southern regions was driven by the local high probability of occurrence of deep-sea species such as blackmouth catshark, velvet belly lanternshark, and rabbitfish (Appendix S11). Although the overall level of uncertainty in the mean predicted species richness was relatively low (SD < 0.4), it exhibited higher values in the central and southern offshore areas, suggesting greater temporal variations (Fig. 1b).

**Figure 1.**
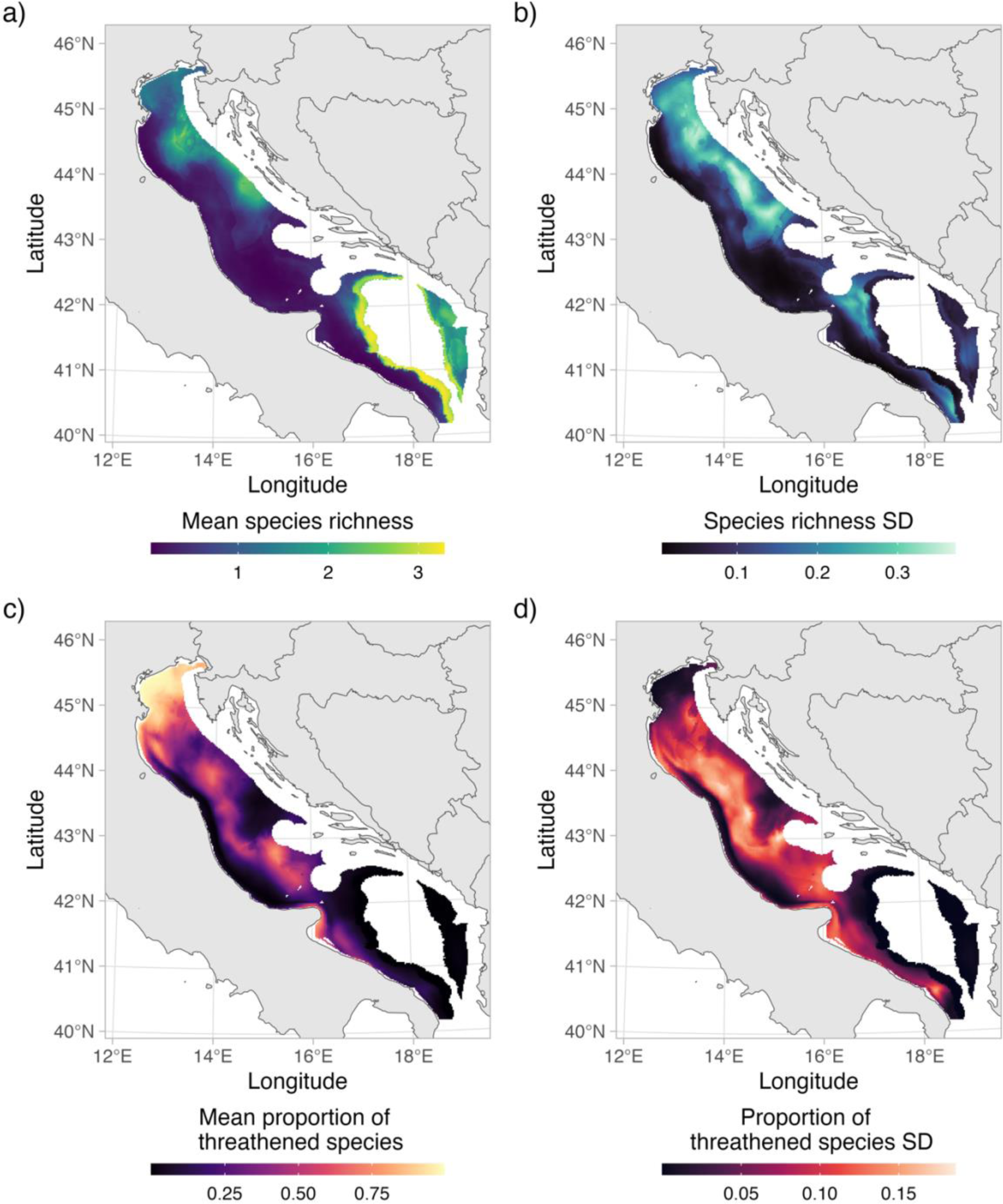
Mean and standard deviation (SD) of species richness (a-b). Mean and SD of proportion of threatened species (c-d). Metrics are computed over the period 2009 to 2021.

The proportion of threatened species was found to be highest in the northernmost part of the study area and lowest along the central and southernmost coasts (Fig. 1c). This pattern was influenced by the relatively high local probability of occurrence of spurdog, smooth-hound, and common eagle ray in the northern regions, all of which are classified as threatened species (Appendix S10). The overall level of uncertainty in the mean proportion of threatened species remained relatively low (SD < 0.2), exhibiting lower values in the northern and central coastal regions as well as in the southern and southeastern offshore areas. This suggests that these specific areas experienced lower temporal fluctuations (Fig. 1d).

### Spatial Overlap Index

For the threatened species, the analysis of the Spatial Overlap Index (SOI) between species and fishing effort across the most recent years (2018-2021) revealed high mean overlap scores for the spurdog, common eagle ray, and smooth-hound in the northernmost part of the study area, both for OTB and TBB (Fig. 2). Additionally, spurdog exhibited a notable level of overlap with OTB along the offshore regions in the northern-central part of the basin. The SOI for the nonthreatened species is reported in Appendix S12-S13. Despite year-to-year fluctuations, the SOI values remained relatively consistent over time, with closer years showing higher similarity (Appendix S14-S17).

**Figure 2.**
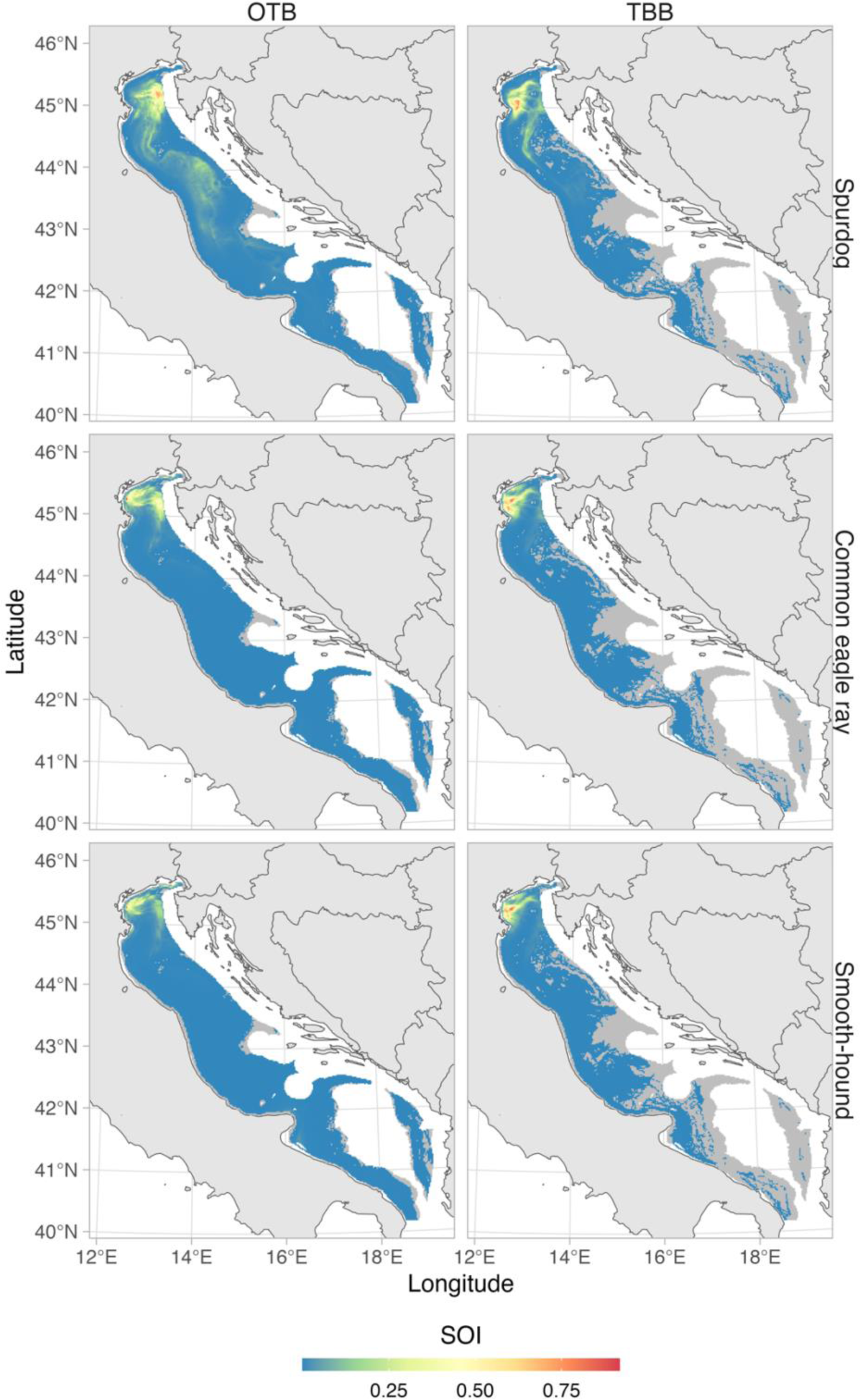
Mean Spatial Overlap Index (SOI) for threatened species and otter bottom trawling (OTB) and beam trawling (TBB) computed over the period 2018-2021. SOI values of 0 are displayed in gray.

### Range overlap

For several species, there was a high potential risk of bycatch extending across a significant portion of their distribution within the study area, as indicated by high range overlap values (RO). Among the species analyzed, starry skate, spurdog, and velvet belly lantern shark were found to share a significant overlap with regions of intensive OTB fishing effort, as indicated by higher RO values (≥ 0.15, averaged across years). Rabbitfish, common eagle ray, blackmouth catshark, smooth-hound, and brown skate exhibited somewhat lower but still significant RO values (< 0.15 and ≥ 0.1, averaged across years) (Fig. 3).

**Figure 3.**
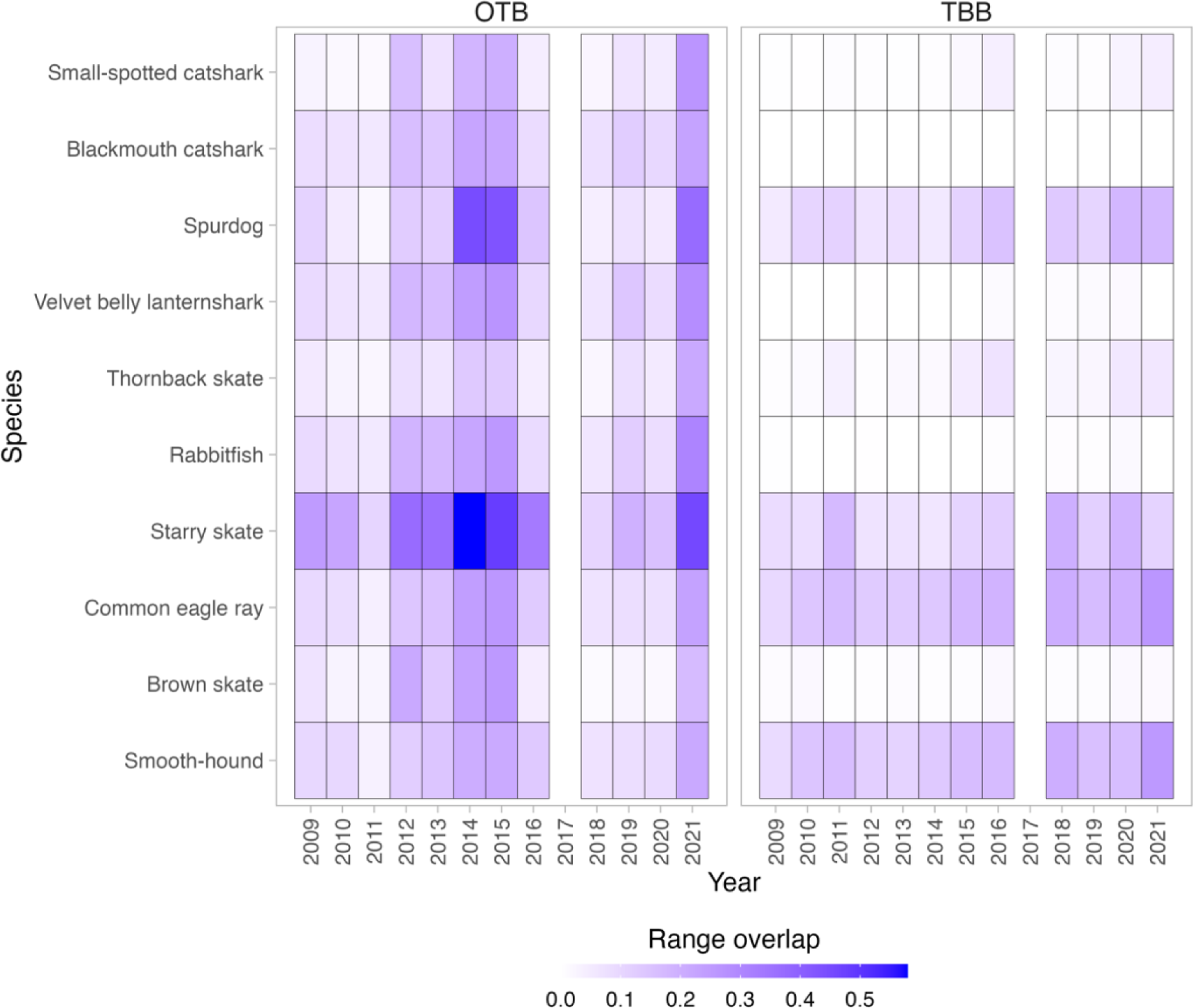
Yearly range overlap (RO) between species and otter bottom trawling (OTB) and beam trawling (TBB) fishing gears. A grid cell is considered to be occupied by a species with ≥ 75th percentile of the probability of occurrence, and by trawling activities with ≥ 75th percentile of fishing effort. The year 2017 is omitted due to sampling occurring outside the designated sampling period (May to September; see the section *Survey data* in *Methods*).

On the other hand, common eagle ray, smooth-hound, spurdog and starry skate had the highest overlap with regions affected by TBB activities (≥ 0.1, averaged across years) (Fig. 3). The RO metrics exhibited variations across years, with significant fluctuations observed for certain species, including spurdog, common eagle ray, and smooth-hound, in both OTB and TBB contexts (Fig. 3). Temporal patterns were also evident, with RO values increasing or decreasing together in certain years, likely in response to changes in fishing effort.These findings remained consistent regardless of the threshold used (Appendix S18-S19).

### Species richness overlap with fishing effort

In general, the areas exhibiting simultaneous high species richness and high OTB fishing effort were predominantly found in the northernmost section of the basin and in some areas along the outer edge of the continental shelf in the southernmost region (Fig. 4). On the other hand, for TBB, our analysis revealed significant overlap between regions of high species richness and fishing effort in the northernmost area of the basin and the northern central region (Fig. 4). Additionally, we identified two distinct zones in the northern-central part of the basin and in the deeper offshore southernmost areas, characterized by high species richness and comparatively low fishing effort of OTB and TBB (Fig. 4).

**Figure 4.**
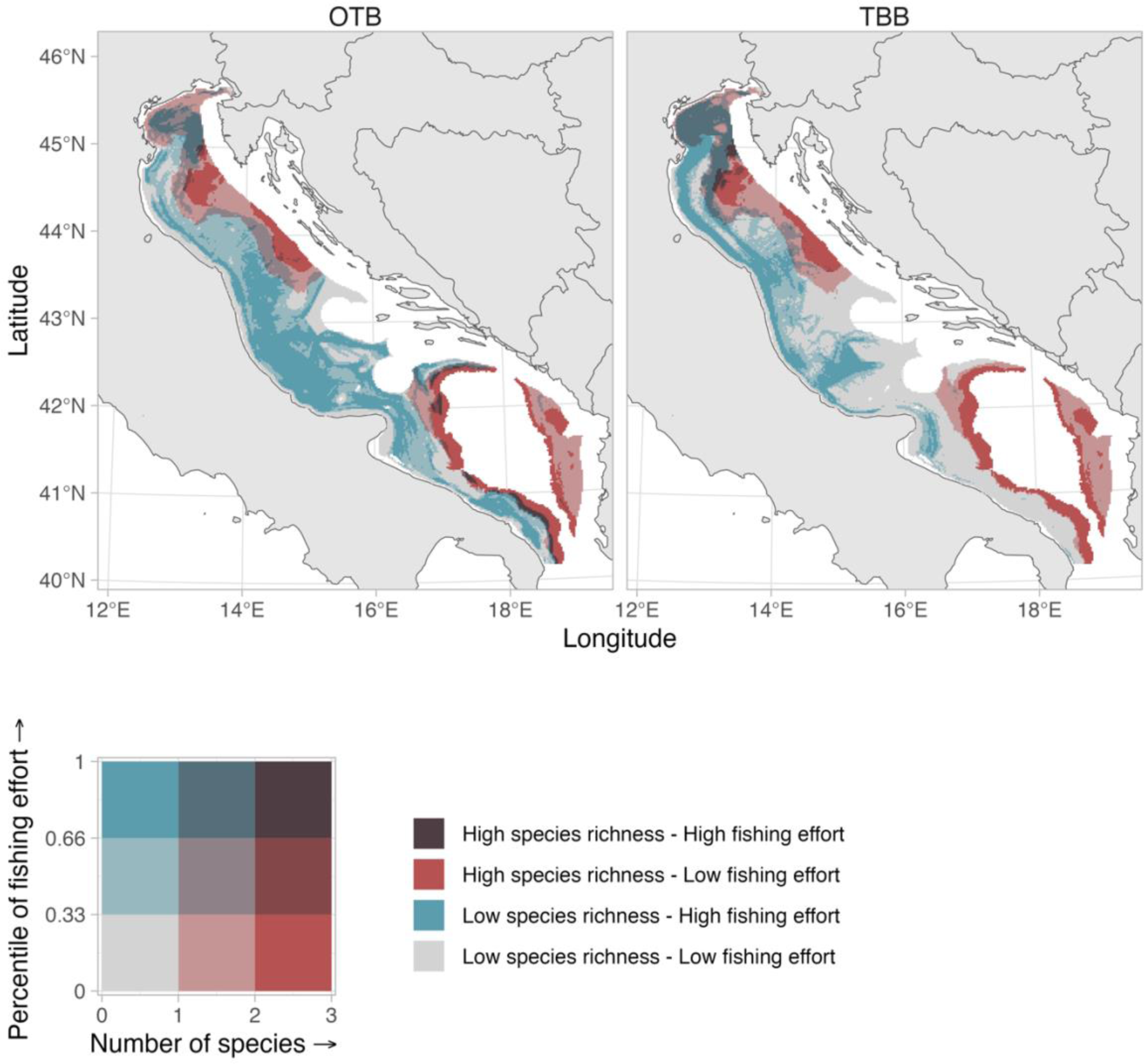
Bivariate plot illustrating the relationship between mean species richness and the percentile of fishing effort for the period 2018-2021 for otter bottom trawling (OTB) and beam trawling (TBB). Dark red areas indicate locations with the highest mean predicted number of species and higest fishing effort. Light red areas correspond to regions characterized by high species richness and low fishing effort. Conversely, dark blue areas represent locations with high fishing effort but a low number of species, whereas gray areas depict regions where both fishing effort and number of species are low.

### Proportion of threatened species overlap with fishing effort

The analysis of OTB revealed distinct patterns in the spatial overlap between fishing effort and the proportion of threatened species. Specifically, in the northernmost regions of the study area, high fishing effort coincided with a higher proportion of threatened species (Fig. 5). However, towards the central and southern portions of the study area, the proportion of threatened species gradually diminishes, in combination with the consistently high levels of OTB fishing activity. Regarding TBB, a clear spatial pattern emerges. The northern part of the study area exhibits a pronounced overlap between fishing effort and the proportion of threatened species. However, this overlap rapidly diminishes southward (Fig. 5). Moreover, we identified distinct areas in both the northernmost and northern-central parts of the basin, which were characterized by a medium-to-high proportion of threatened species and relatively low fishing effort of OTB and TBB (Fig. 5).

**Figure 5.**
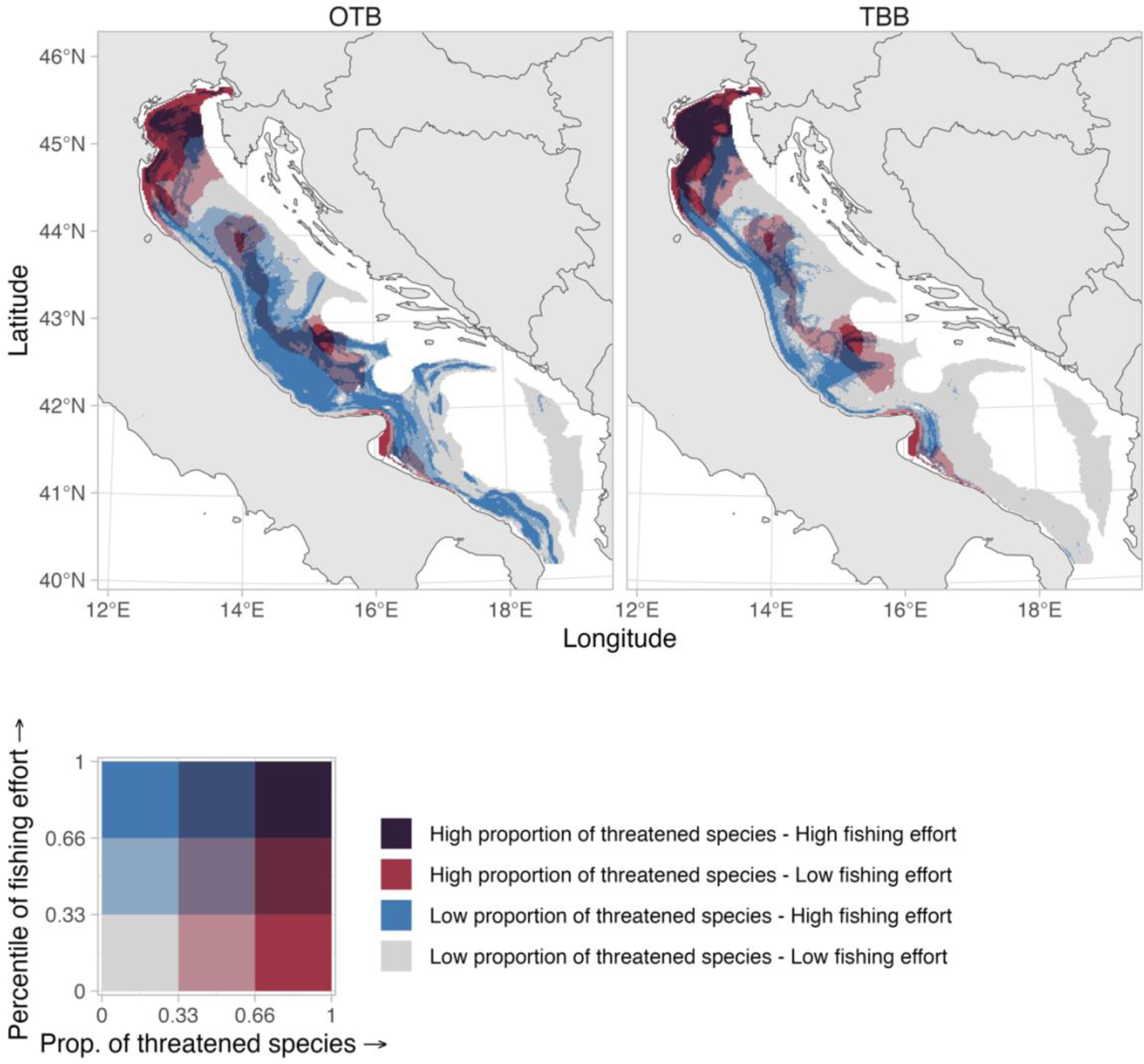
Bivariate plot illustrating the relationship between the mean proportion (prop.) of threatened species and the percentile of fishing effort for the period 2018-2021 for otter bottom trawling (OTB) and beam trawling (TBB). Dark red areas indicate locations with the highest mean predicted proportion of threatened species and fishing effort. Light red areas correspond to regions characterized by a high proportion of threatened species and low fishing effort. Conversely, dark blue areas represent locations with high fishing effort but a low proportion of threatened species, whereas gray areas depict regions where both fishing effort and the proportion of threatened species are low.

## Discussion

Effective marine conservation and fisheries management requires a comprehensive understanding of the spatial overlap between species distribution and fishing activities (Douvere, 2008). Especially for chondrichthyans, the deficiency of such information is often recognized as a key factor contributing to the failure to implement suitable management measures (Cavanagh & Gibson, 2007; FAO, 2022). Bottom trawl fishing is known to result in incidental bycatch of nontarget species, including chondrichthyans (Carpentieri et al., 2021). In addition to the direct removal of chondrichthyans, bottom trawl fishing also poses a threat to their habitats. The use of weighted nets dragged along the seabed can cause disturbance and destruction of benthic habitats and removal of the associated biota (ICES, 2023; Sciberras et al., 2018; Watling & Norse, 1998), which are crucial for the reproduction, sheleter and foraging of many chondrichthyans (e.g. Carrier & Pratt, 1998; Heithaus et al., 2002; Simpfendorfer & Milward, 1993).

In this study, we propose an innovative approach that produces model-based overlap between chondrichthyan distribution and bottom trawl fishing, leveraging high-resolution Vessel Monitoring System (VMS) and the spatial distribution of species derived from geostatistical models fitted to fishery-independent survey data. This information may serve as a critical tool for identifying areas that require protection and implementing conservation measures, such as establishing protected zones or imposing fishing restrictions. Such measures can play a crucial role in safeguarding vulnerable chondrichthyan species and communities, thus promoting their conservation and recovery. We applied this integrated approach to the Adriatic Sea, an highly exploited area (Amoroso et al., 2018) characterized by a long history of chondrichthyan depletion (Ferretti et al., 2013) and limited data on their incidental catches (FAO, 2022). We addressed the latter knowledge gap providing novel insights into the potential interaction between chondrichthyans and bottom trawl fishing in this basin.

Our findings reveal significant spatial overlap between chondrichthyans and bottom trawl fishing hotspots in the western part of the Adriatic Sea. This overlap raises concerns about the vulnerability of chondrichthyans, that also includes threatened shark and ray species, to fishing pressure, particularly in regions with high fishing intensity. Our results show distinct spatial patterns of overlap, highlighting both areas of concern and potential refuge for chondrichthyan populations.

We detected two distinct species richness hotspots within the study domain. One hotspot, situated in the northern and central offshore region of the basin, featured primarily threatened species like spurdog, smooth-hound, and common eagle ray, as well as some non-threatened species including thornback skate and small-spotted catshark. The second hotspot, located in the southern and deepest regions, was characterized by the presence of non-threatened deep sea species such as blackmouth catshark, velvet belly lanternshark, and rabbitfish.

In the northernmost part of the Adriatic Sea, we observed a pronounced overlap between the distribution of a high proportion of threatened chondrichthyan species (i.e. spurdog, smooth-hound, and common eagle ray) and hotspots of bottom trawl fishing activity. Moreover, for these species, this significant spatial overlap extended over a considerable portion of their habitat, despite some year-to-year fluctuations. These areas of high spatial overlap suggest potential risks for threatened chondrichthyan populations. Conversely, we also identified areas with minimal spatial overlap between high chondrichthyans probability of occurrence and bottom trawl fishing, indicating potential refuge areas. One such area, known as “area dei fondi sporchi” (or “dirty area”) by local fishers, located in the northern-central part of the basin, contains relict sand rich in epifaunal organisms that make trawling difficult or even impossible (Scaccini, 1967).

Furthermore, our analysis revealed localized differences in spatial overlap between different fishing gears, such as otter bottom trawling (OTB) and beam trawling (TBB), along the central and southern Adriatic coastline. These differences highlight the significance of taking specific fishing gears, and their respective potential impacts, into account when evaluating spatial overlap and formulating effective management strategies. Notably, both OTB and TBB are the focal gears addressed within the Multiannual Management Plan (GFCM, 2019) for demersal species in the Adriatic Sea including hake (*Merluccius merluccius*), red mullet (*Mullus barbatus*), common cuttlefish (*Sepia oficinalis*), Norway lobster (*Nephrops norvegicus*) and common sole (*Solea solea*) (Sea Around Us, 2006). This Management Plan combines different measures and among them, fishing effort regimes and spatio-temporal closures. Tailoring conservation measures, within this management context, related to these fishing gears and their interactions with chondrichthyan populations, can optimize conservation efforts and minimize unintended bycatch or habitat damage.

In addition to identifying hotspots of overlap between species or proportion of threatened species with fishing, our model-based approach also allows calculation of range overlaps. This is an important metric along identifying hotspots of overlap, because together with distribution models, it provides information about potential effects of relocating spatial fishing effort. For instance, for species with clear spatial hotspots and low range overlap, relocating fishing effort might reduce mortality from incidental bycatch substantially within the domain. However, for species with high range overlap, simply limiting fishing in “the hottest spot” might have neglible effects, if the fishery over the entire species range and the area with high overlap is large. This illustrates the need to develop a suite of indices before implementing spatial fisheries-restrictions. Understanding where fishing would increase after such an intervention, and how that affects the total mortality rates from fishing, is one of the main challenges in designing protected areas successfully (Hilborn et al., 2004; Ovando et al., 2021).

While our study provides novel insights into the spatial overlap between chondrichthyan distribution and bottom trawl fishing in the western part of the Adriatic Sea, there are certain limitations that should be considered when interpreting the results. Firstly, our analysis relied on the availability and quality of data sources, which may introduce uncertainties and biases. For instance, VMS data is considered a good proxy for fishing effort for the fleet segment larger than 15 meters in overall length, while their reliability decreases for vessels between 12 and 15 meters in length and miss the fishing effort of the fleet below 12 meters. This may result in only a partial representation of fishing activities, potentially leading to an incomplete understanding of the spatial overlap between fishing and chondrichthyan distribution for smaller vessels. Secondly, our data only cover the Italian fleet, while other fishing countries can operate outside the Italian territorial sea. Nonetheless, approximately 70% of the fishing fleet operating in the Adriatic Sea belongs to Italy (FAO, 2022) and therefore we cover the majority of the fleet that should have the major impact on the chondrichthyans in the study area. Additionally, SDMs are subject to inherent limitations, such as uncertainties in species occurrence data, model assumptions, and potential omission of relevant environmental variables. However, the approach described here, calculating overlap on model-predicted probabilities of occurrence also has benefits, primarily in that it is not as sensitive to temporal changes in sampling across in space (Thorson et al., 2016). Lastly, the spatial resolution of our analysis may influence the observed patterns of overlap. While we aimed to use the finest available resolution of the data, the accuracy and precision of VMS data and species occurrence records can vary spatially. Fine-scale variations in both fishing effort and chondrichthyan distribution may not have been fully captured in our analysis, potentially leading to underestimation or overestimation of the true spatial overlap. Despite these mentioned contraints, the approach presented here, to the best of our knowledge, offers the most comprehensive information available regarding the overlap of fishing effort with the chondrichthyan community in the Adriatic Sea.

Our analysis focused on a specific period of the year (from the late spring to the end of summer) due to the lack of consistent autumn and winter fisheries-independent surveys during the last twenty years. Seasonal variations in fishing effort and chondrichthyan distribution may occur though, and therefore caution must be taken in extrapolating our results to other seasons, especially for conservation and management purposes.

While spatial overlap between fishing activities and chondrichthyan distribution indicates potential interaction, it does not directly provide evidence of actual biological impacts or quantify the extent of such impacts on chondrichthyan populations. Nevertheless, Mediterranean chondrichthyans are widely acknowledged to be highly vulnerable to trawlers (Carpentieri et al., 2021; Cavanagh & Gibson, 2007; FAO, 2022). Given the limited availability of direct and quantitative official bycatch data, our results may offer an initial and valid approach to inform conservation efforts and guide precautionary management decisions.

In conclusion, our study introduces an innovative approach that combines VMS data with geostatistical species distribution models, allowing us to analyze the spatial co-occurrence of chondrichthyan species and commercial bottom trawl fishing practices in the Adriatic Sea. Through this analysis, we identified areas of heightened risk for bycatch and thus of potential impacts on chondrichthyan populations, as well as areas that could serve as potential refuge. These findings provide therefore new knowledge that may support the implementation of targeted conservation efforts and the development of effective spatial management measures in the region.

## Acknowledgments

We thank the MEDITS staff involved in the scientific sampling and analysis of biological data. This study has been conceived under the International PhD Program “Innovative Technologies and Sustainable Use of Mediterranean Sea Fishery and Biological Resources” (http://www.fishmed-phd.org/).

