## Supporting Information for "Assessing the overlap between fishing activities and chondrichthyans distribution exposes high-risk areas for bycatch of threatened species"

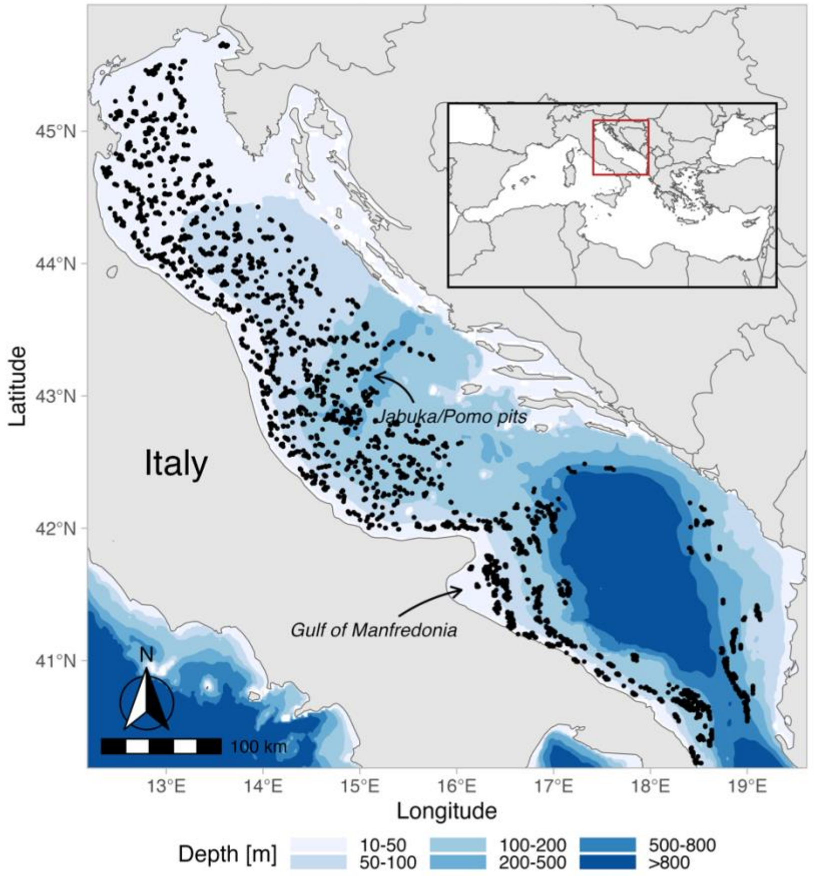


Appendix S1. Location of the study area and the positions of MEDITS bottom trawl hauls swept during the period 1999–2021 selected and analyzed in this study.

Appendix S2. Environmental covariates included in the best fitting model for each species, along with their respective explanatory power. Species are ordered according to model selection similarities. Green cells represent the combinations of covariates tested during model selection, while the '+' sign indicates the terms that were retained by the best fitting model, as selected by Akaike Information Criterion. T = temperature; Oxy = oxygen; T:Oxy = interaction term between T and Oxy; AUC = Area Under the Curve.

| Species | Depth | log(Depth) | Depth + Depth$^2$ | log(Depth) + log(Depth$^2$) | T | T + T$^2$ | Substrate | Oxy | T:Oxy | Conditional R2 | AUC |
|---|---|---|---|---|---|---|---|---|---|---|---|
| Spurdog | | | | + | | + | + | + | + | 0,89 | 0,98 |
| Small-spotted catshark | | | + | | + | | + | + | + | 0,84 | 0,97 |
| Thornback skate | | | | + | | + | + | + | | 0,76 | 0,96 |
| Blackmouth catshark | | | + | | | | | | | 0,86 | 0,99 |
| Velvet belly lanternshark | | | + | | | | | | | 0,92 | 0,99 |
| Rabbit fish | | | + | | | | | | | 0,88 | 0,99 |
| Starry skate | | | | + | | + | | | | 0,54 | 0,91 |
| Common eagle ray | | | + | | + | | | | | 0,86 | 0,98 |
| Smooth-hound | | | | + | + | | | | | 0,76 | 0,98 |
| Brown skate | | | | + | | | | | | 0,92 | 0,98 |

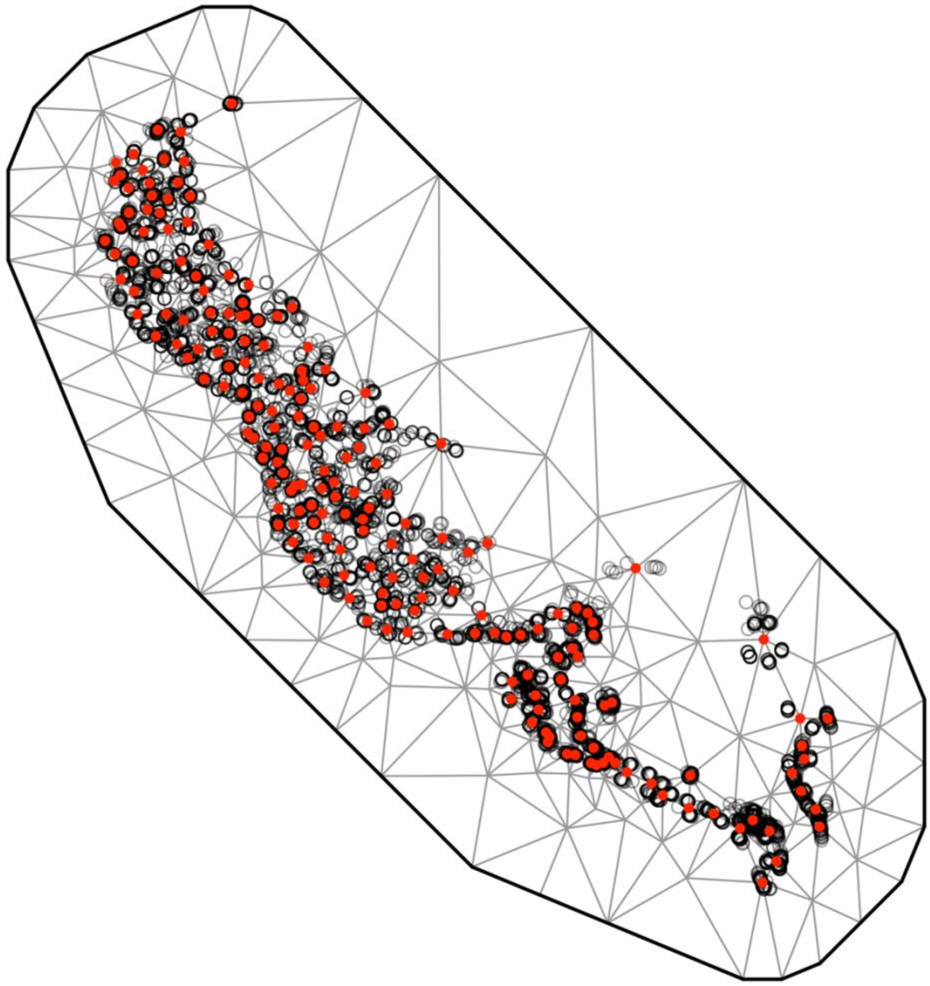

Appendix S3. Stochastic Partial Differential Equation (SPDE) mesh for the species distribution models. The knots (n=200) are displayed in red, and the haul positions are represented as open circles.

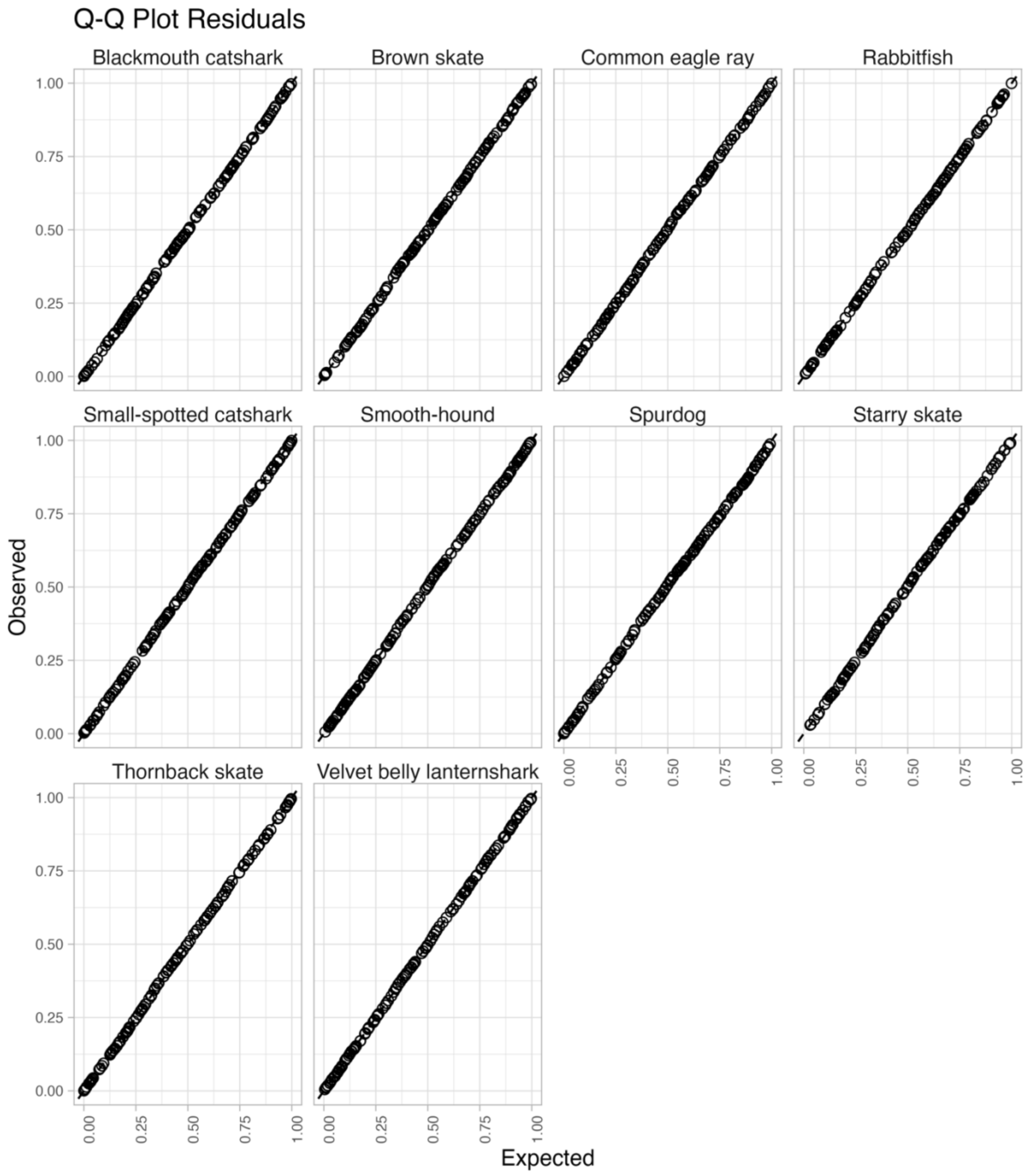


Appendix S4. Quantile-quantile plot illustrating randomized quantile residuals of the species distribution models. To enhance clarity, a random sample of 1000 points was selected and visualized.

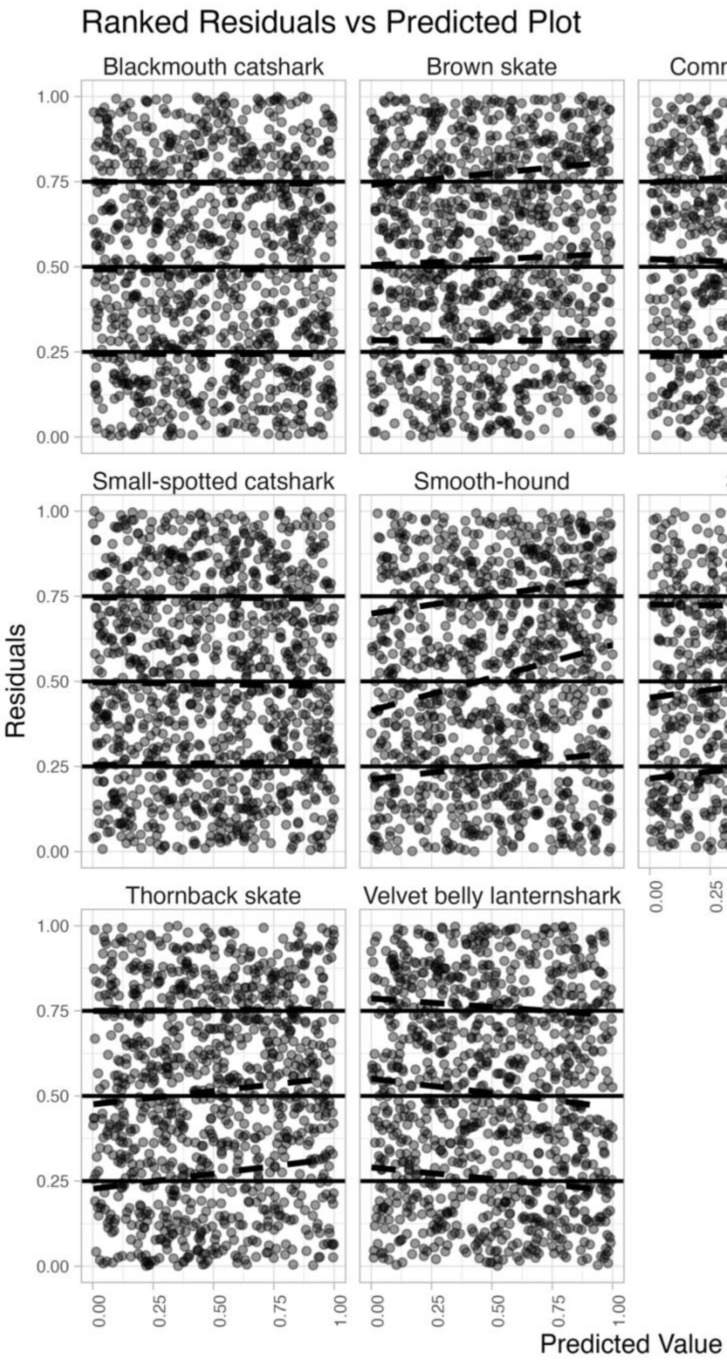


Appendix S5. Quantile regression of the residuals of the species distribution models. Solid lines represent the theoretical expectations for the 0.25, 0.5, and 0.75 quantiles, while dashed lines represent the observed values. To enhance clarity, a random sample of 1000 data points was selected and displayed.

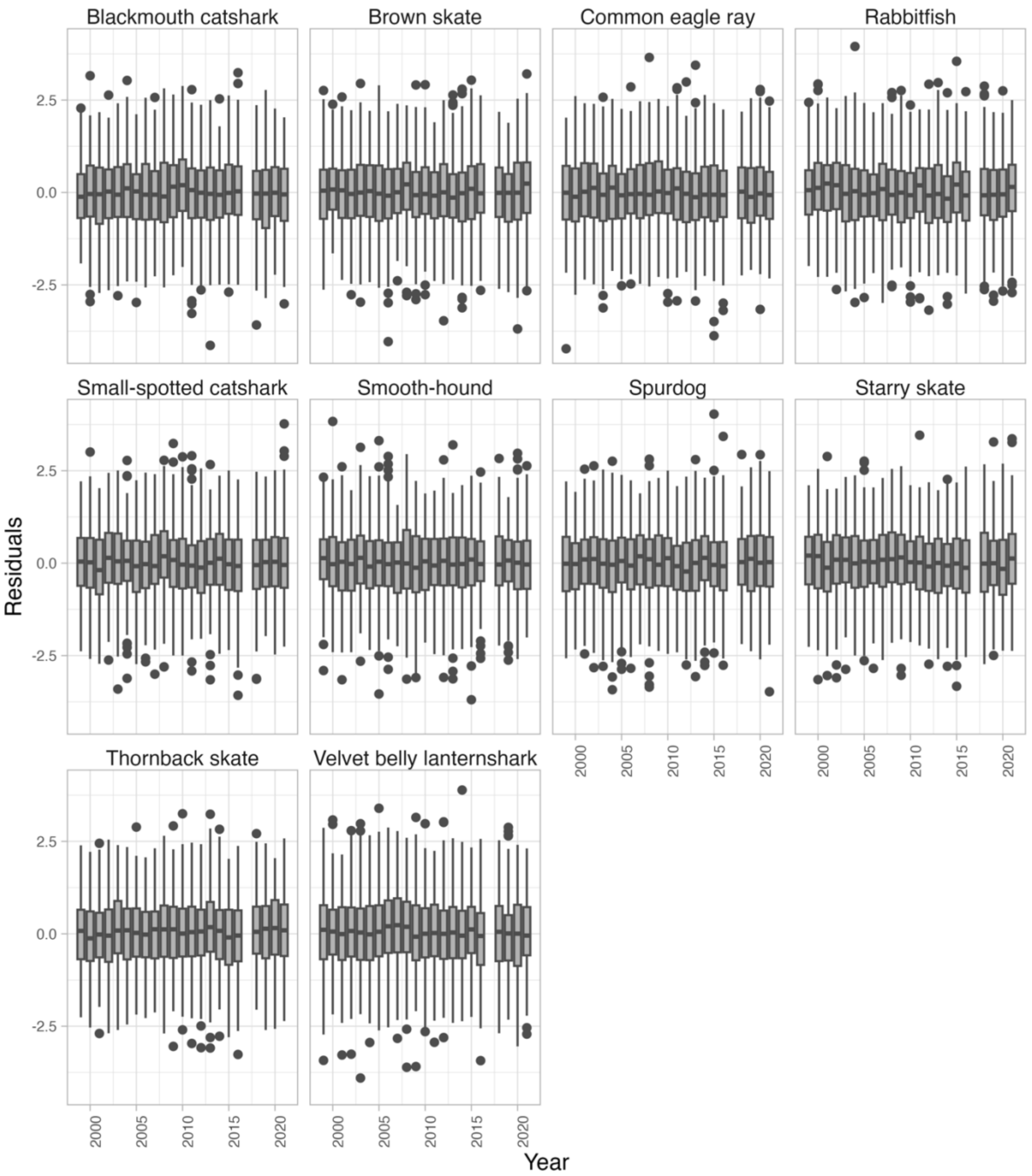


Appendix S6. Residuals of the species distribution models aggregated by year. The year 2017 is omitted due to sampling occurring outside the designated sampling period (May to September; see the section *Survey data* in *Methods*).

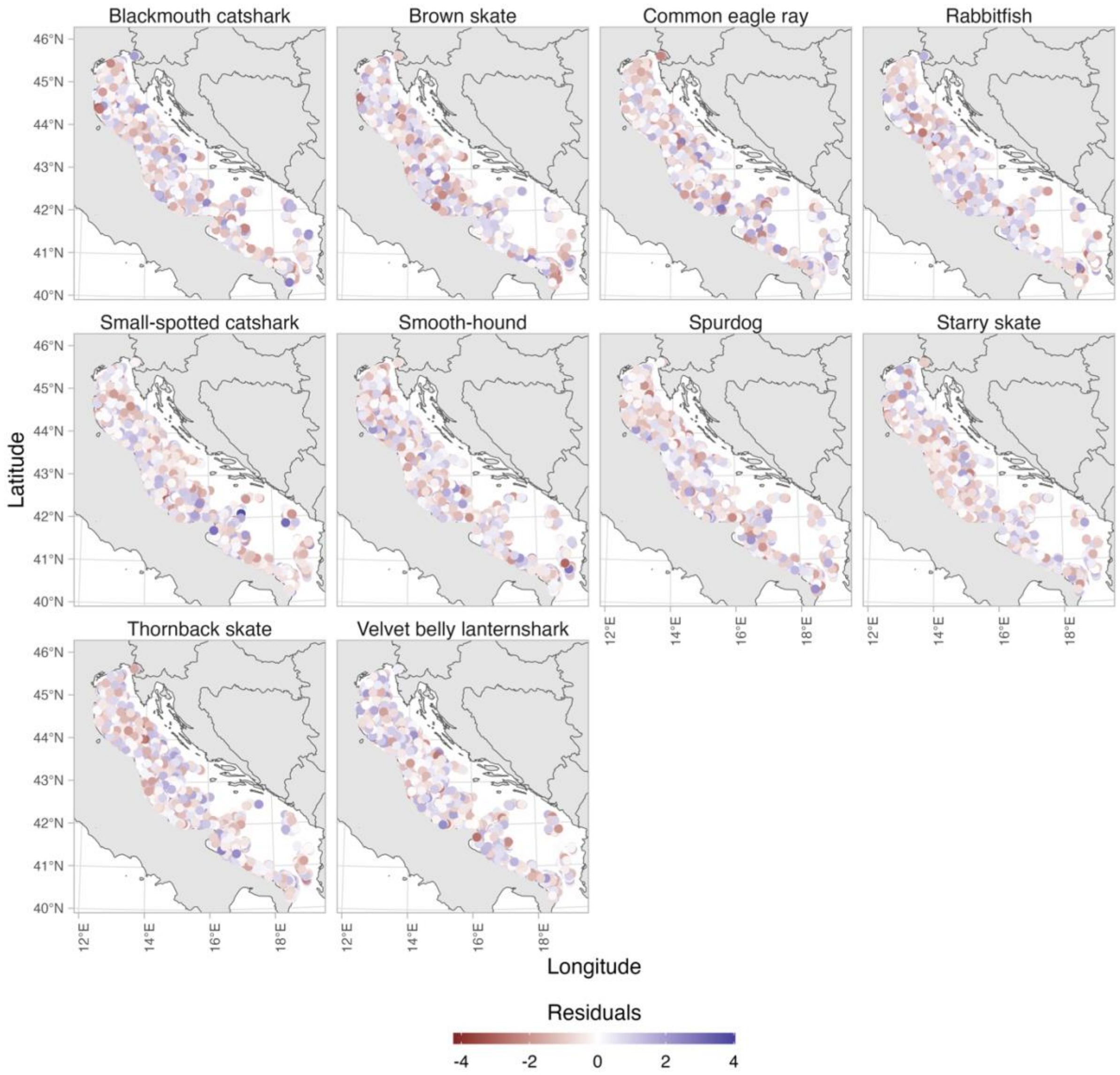


Appendix S7. Residuals of the species distribution models plotted in space.

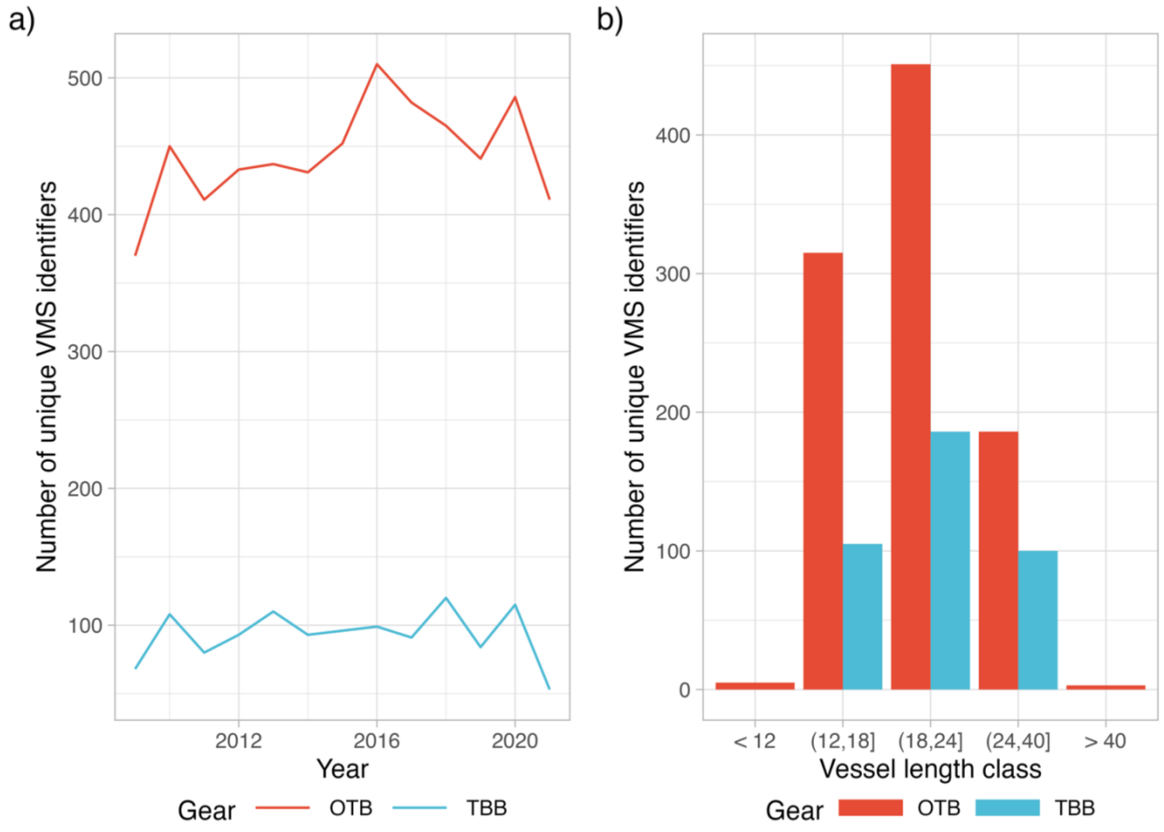


Appendix S8. Number of unique VMS identifiers per year separated into otter bottom trawling (OTB) and beam trawling (TBB) (a). Number of unique VMS identifiers categorized by vessel length class (b).

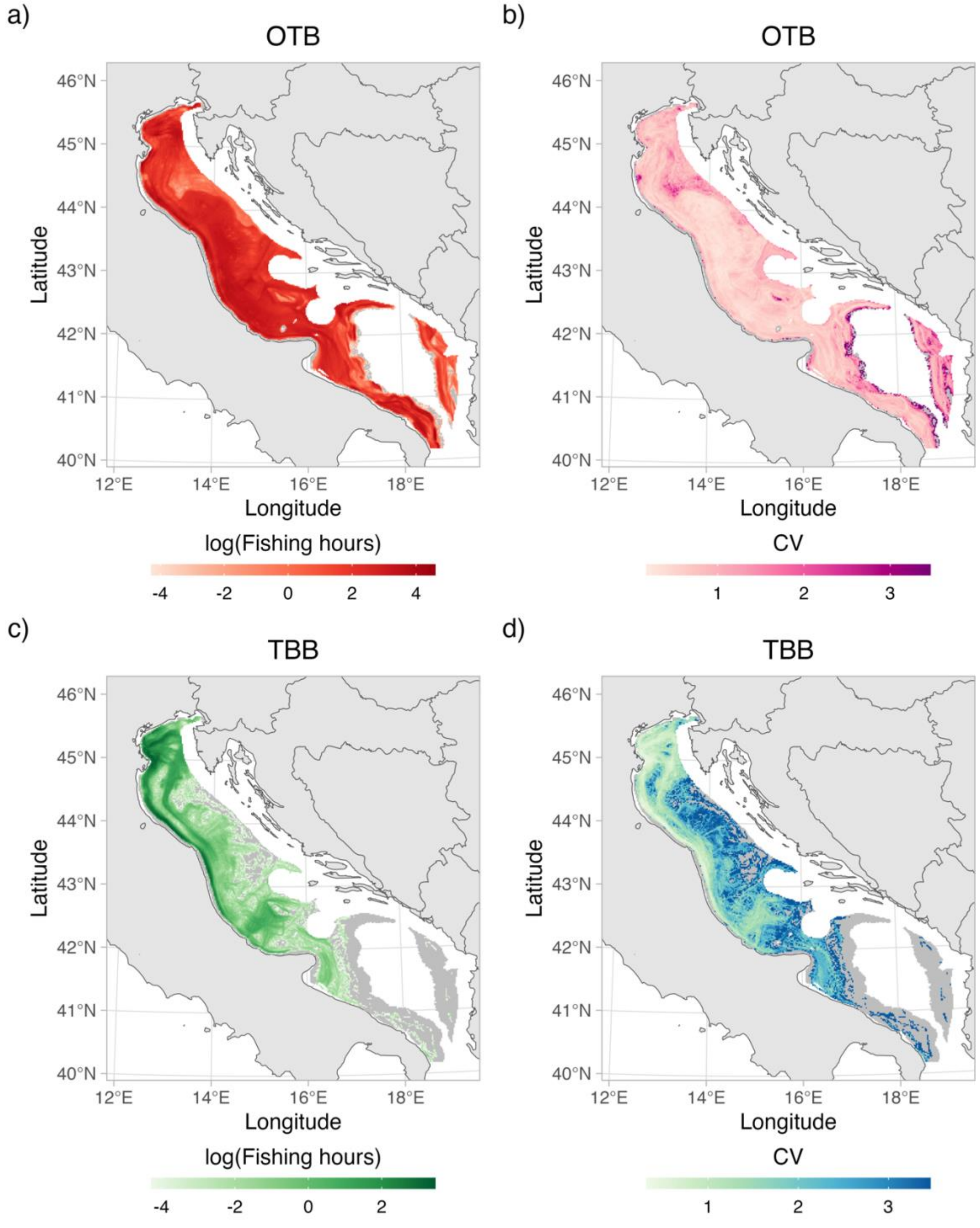


Appendix S9. Mean fishing effort (fishing hours) derived from VMS signals. Mean fishing effort for otter bottom trawlers (OTB) (a) and beam trawlers (TBB) (c). Coefficient of variation (CV) of fishing effort for OTB (b) and TBB (d). Gray areas represent regions with estimated 0 fishing effort. The plots represent the sum of effort from May to September averaged over the period 2009 to 2021.

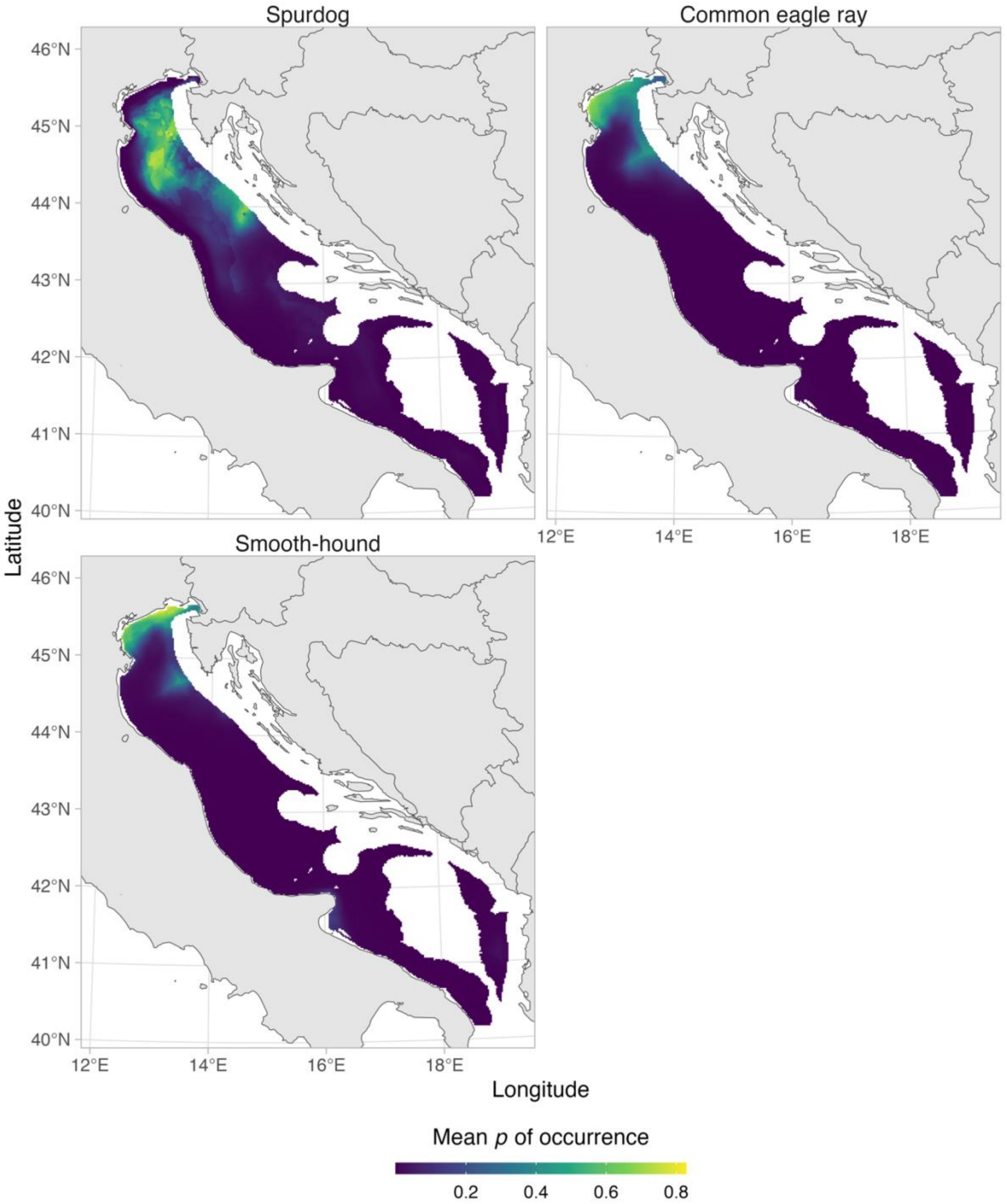


Appendix S10. Mean predicted probability (*p*) of occurrence for the threatened species. Means are computed over the period 2009 to 2021.

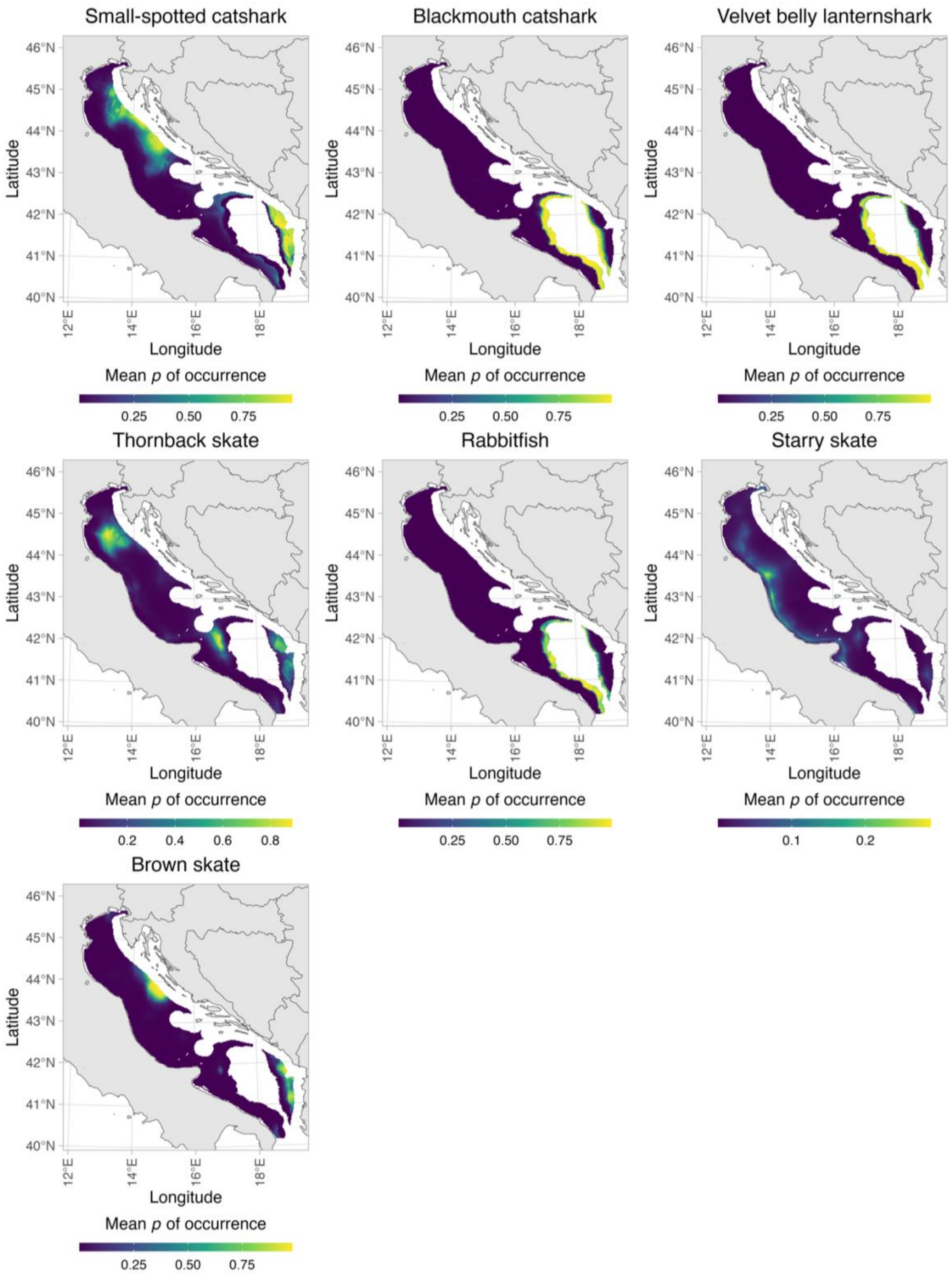


Appendix S11. Mean probability ($p$) of occurrence for the nonthreatened species. Means are computed over the period 2009 to 2021.

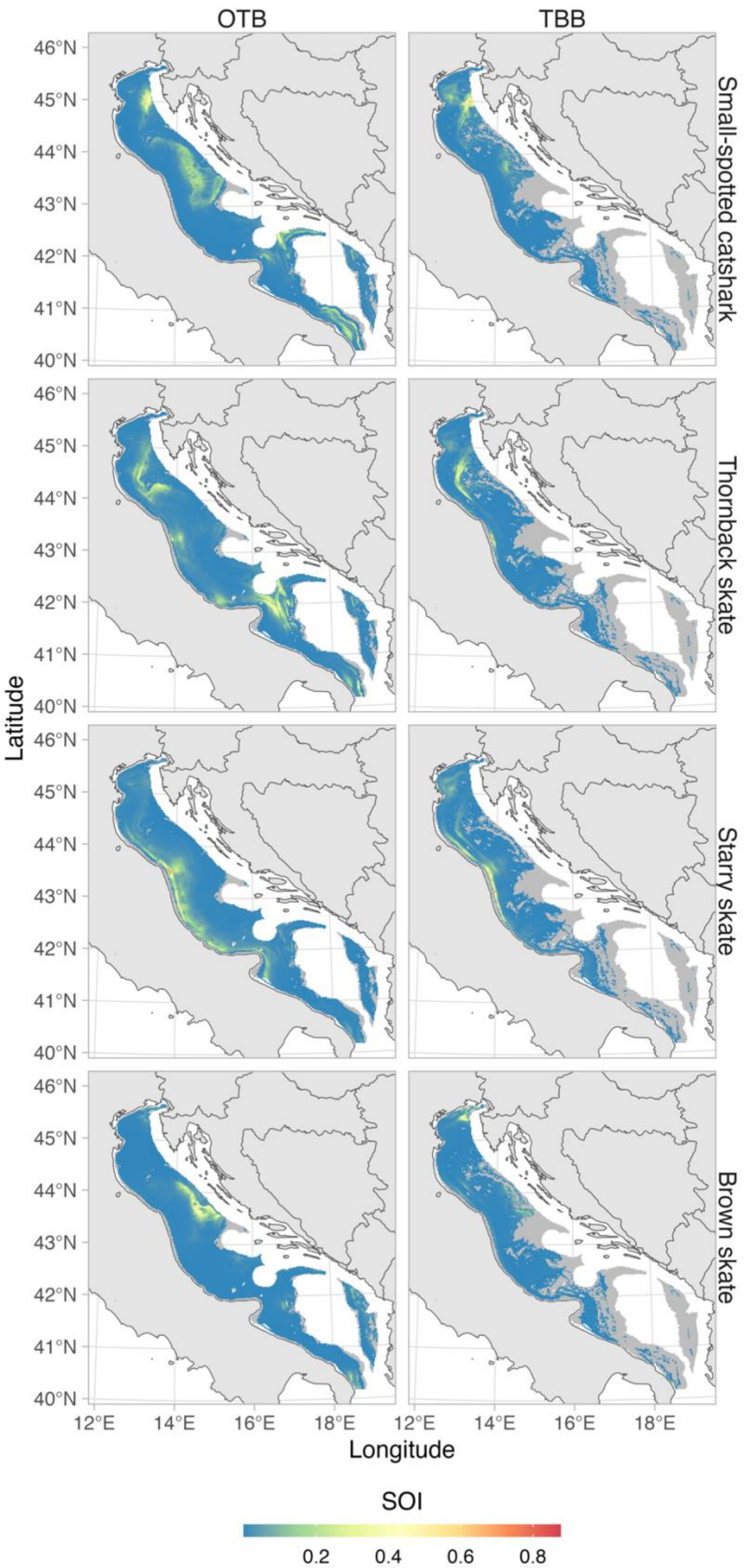


Appendix S12. Mean Spatial Overlap Index (SOI) for nonthreatened species computed over the period 2018-2021. SOI values of 0 are displayed in gray.

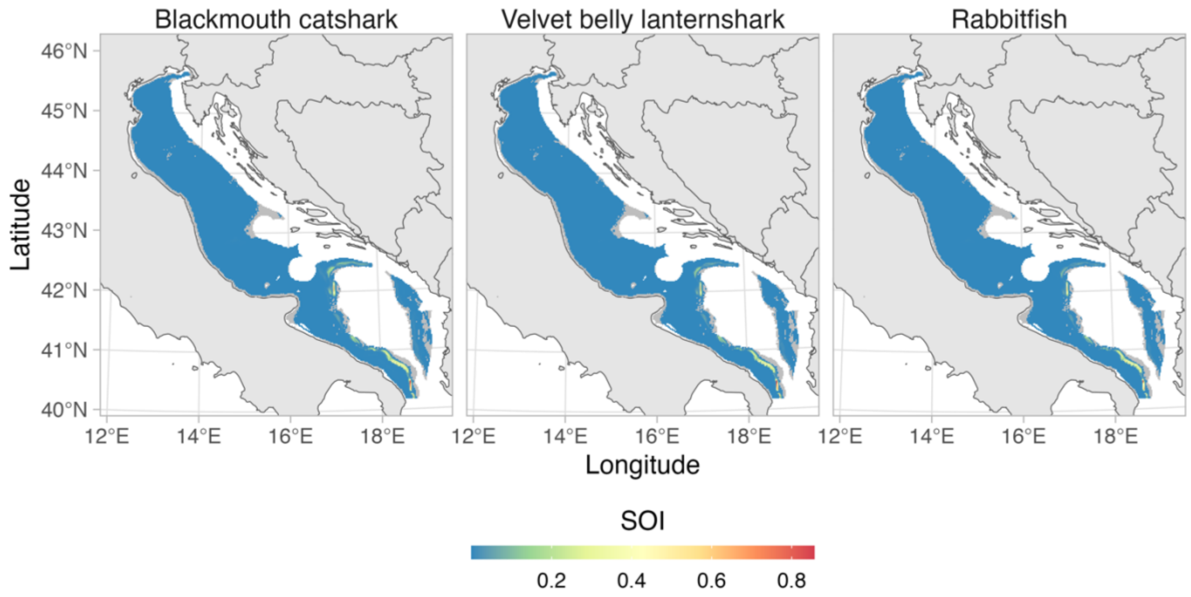


Appendix S13. Mean Spatial Overlap Index (SOI) between for the remaining nonthreatened species and OTB for the years 2018-2021. SOI values of 0 are displayed in gray. For simplicity, only SOI with OTB is shown due to virtually 0 overlap of these deep-sea species with TBB.

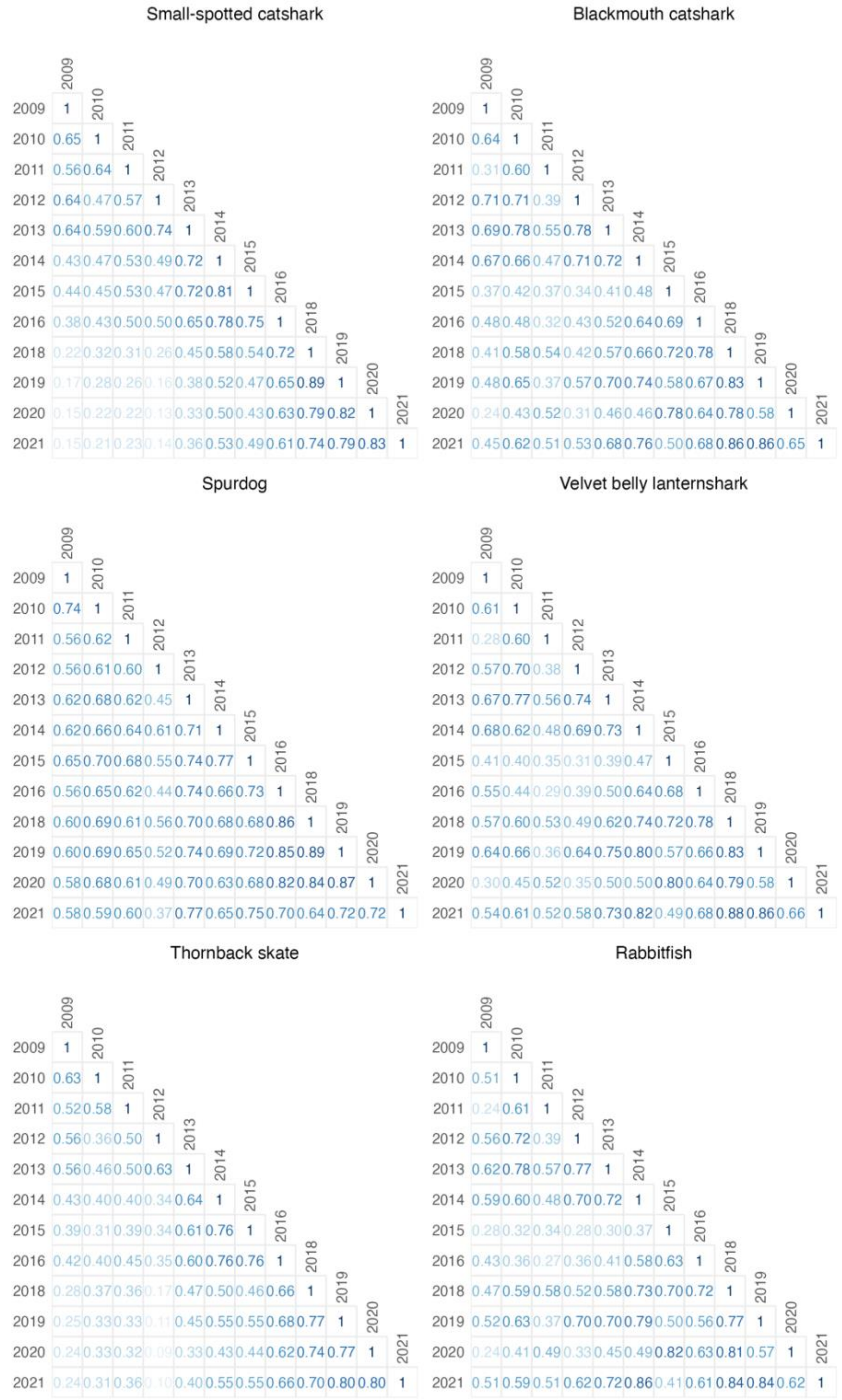


Appendix S14. Inter-year correlations of Spatial Overlap Index for some species and OTB. Values in blue inside the plot represent Pearson correlation coefficients.

.

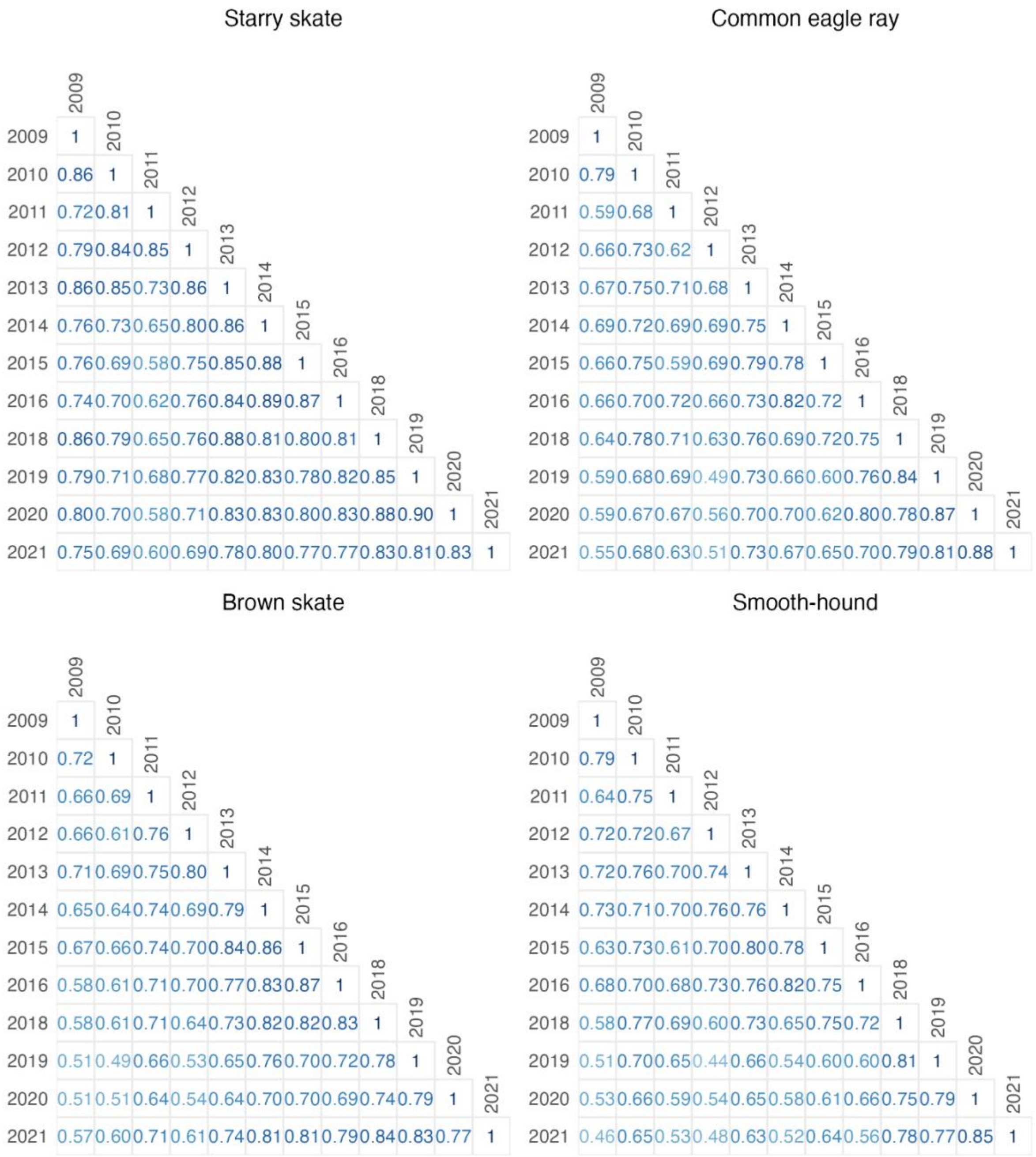


Appendix S15. Inter-year correlations of Spatial Overlap Index for the remaining species and OTB. Values in blue inside the plot represent Pearson correlation coefficients.

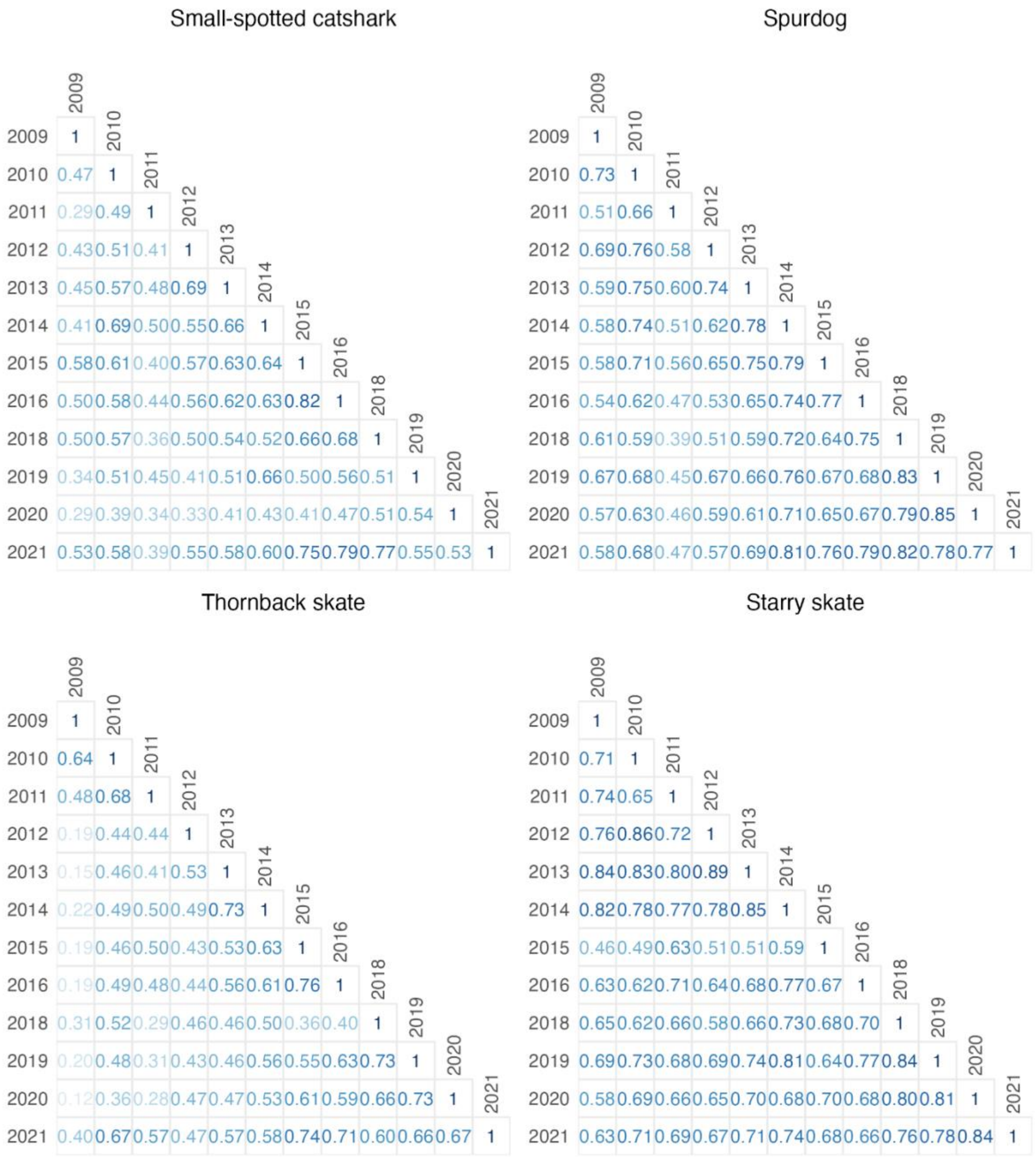


Appendix S16. Inter-year correlations of Spatial Overlap Index for some species and TBB. Values in blue inside the plot represent Pearson correlation coefficients.

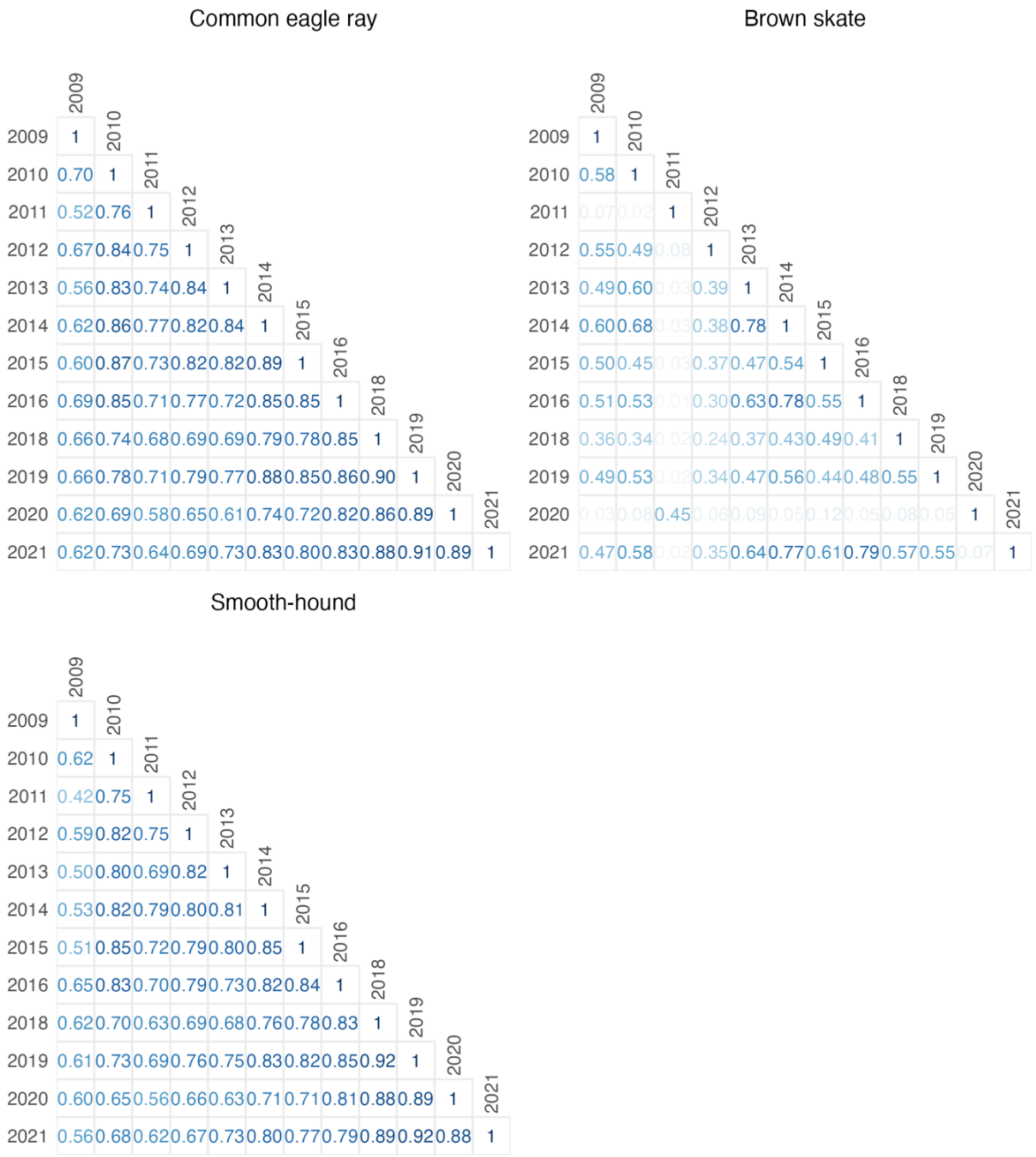


Appendix S17. Inter-year correlations of Spatial Overlap Index for the remaining species and TBB. Inter-year correlations of deep-sea species (blackmouth catshark, velvet belly lanternshark and rabbitfish) and TBB are not shown as their overlap is virtually 0. Values in blue inside the plot represent Pearson correlation coefficients.

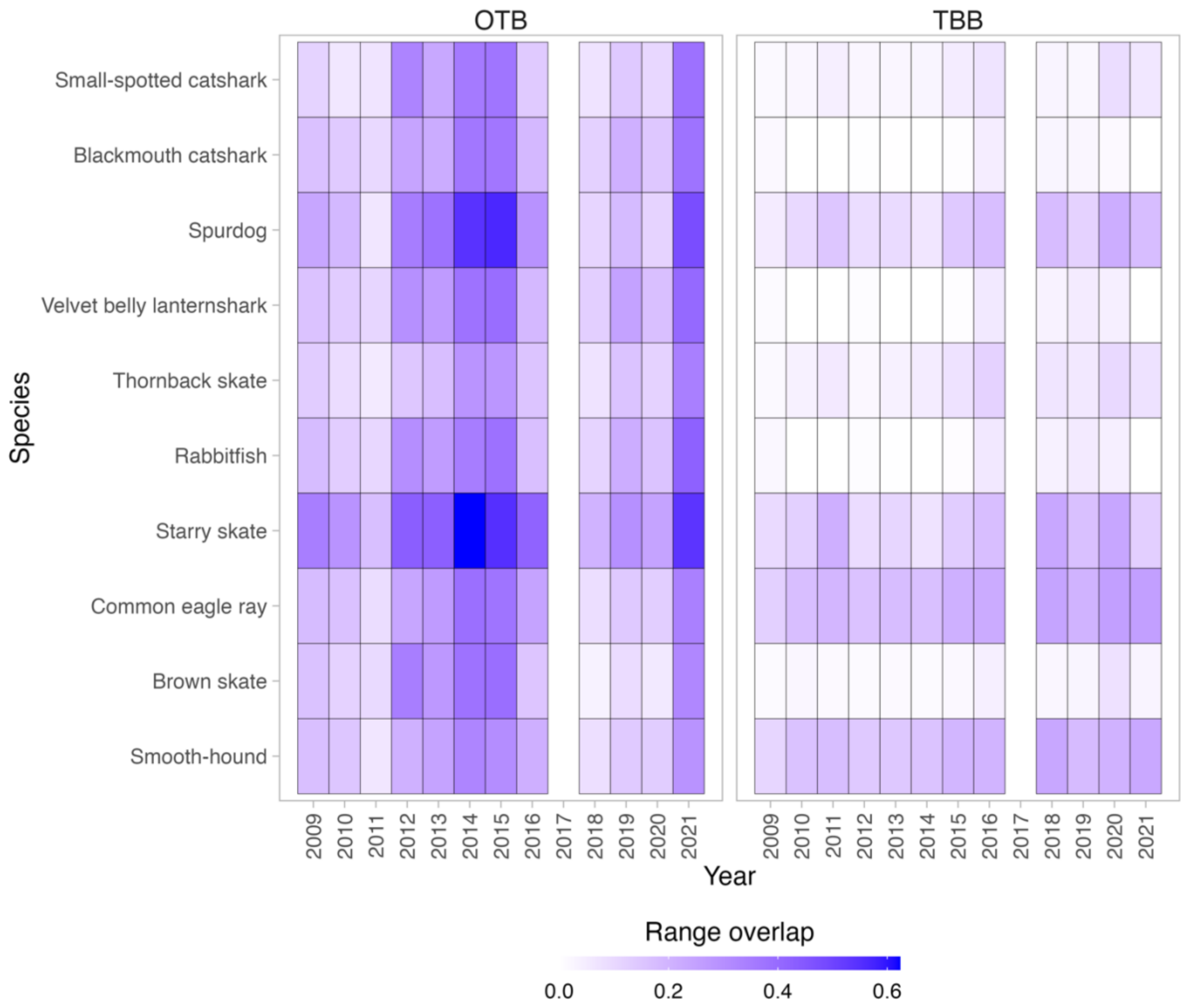


Appendix S18. Yearly range overlap between species and otter bottom trawling (OTB) and beam trawling (TBB) fishing gears. The analysis considered a grid cell to be occupied by a species if it exceeded the 65th percentile of the probability of occurrence, and by trawling activities if it exceeded the 65th percentile of fishing effort. The year 2017 is omitted due to sampling occurring outside the designated sampling period (May to September; see the section *Survey data* in *Methods*).

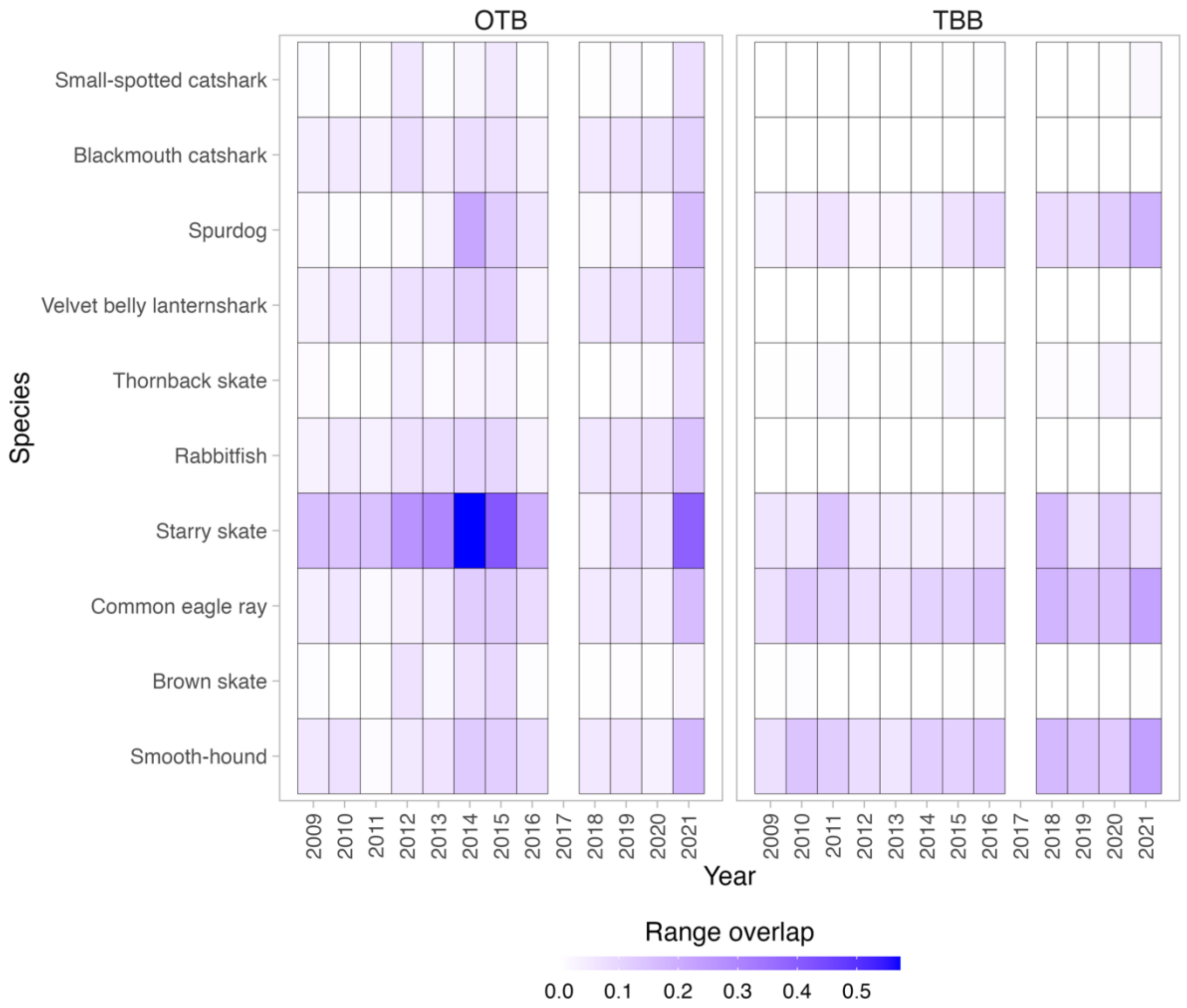


Appendix S19. Yearly range overlap between species and otter bottom trawling (OTB) and beam trawling (TBB) fishing gears. The analysis considered a grid cell to be occupied by a species if it exceeded the 85th percentile of the probability of occurrence, and by trawling activities if it exceeded the 85th percentile of fishing effort. The year 2017 is omitted due to sampling occurring outside the designated sampling period (May to September; see the section *Survey data* in *Methods*).
